# The transcription factor IRF-5 is essential for the metabolic rewiring of CD8 T cells during chronic infection

**DOI:** 10.1101/2024.01.29.577789

**Authors:** Linh Thuy Mai, Sharada Swaminathan, Trieu Hai Nguyen, Tania Charpentier, Hamza Loucif, Liseth Carmona-Pérez, Alain Lamarre, Krista M. Heinonen, Jörg H. Fritz, Simona Stäger

**Author notes:** contributed equally (Sharada Swaminathan, Trieu Hai Nguyen). Address correspondence to: Simona Stäger INRS – Centre Armand-Frappier Santé Biotechnologie 531 Boulevard des Prairies Laval (QC), H7V 1B7, Canada.

## Abstract

Numerous transcription factors are involved in promoting an intricate gene expression program that leads to CD8 T cell exhaustion. Here, we found that the transcription factor IRF-5 is involved in limiting functional exhaustion of CD8 T cells by regulating the cell cycle and contributing to sustaining the mitochondrial functions and oxidative phosphorylation during the chronic stage of LCMV Cl13 infection. CD8 T cells lacking IRF-5 display reduced survival capacity and show increased signs of functional exhaustion during the chronic stage of infection. IRF-5-deficiency also resulted in a severely defective lipid metabolism, in a faulty mitochondrial envelope, and in the reduced capacity to produce ATP. Additionally, we observed increased lipid peroxidation in CD8 T cells lacking IRF-5, when compared with WT cells. These findings identify IRF-5 as a pivotal regulator of the metabolic rewiring that occurs in CD8 T cells during the chronic stages of infection and highlight its role in protecting cells from cell death, possibly by lipid peroxidation.

**Summary:** IRF-5 is critical for regulating mitochondrial functions and oxidative phosphorylation in CD8 T cells during chronic stages of LCMV Cl13 infection.

## Introduction

Maintenance of CD8 T cells is essential for controlling infections caused by intracellular pathogens. However, CD8 T cell responses gradually fade during the chronic stages of infection and antigen specific CD8 T cells become exhausted as a result of overwhelming antigenic stimulation (1–3). Exhausted T cells (Tex) are characterized by an impaired proliferation, the progressive loss of effector functions, the elevated expression of inhibitory receptors, and an altered cellular metabolic profile (4). Tex initially lose their capacity to produce interleukin 2 (IL-2), followed by tumor necrosis factor (TNF) and ultimately interferon gamma (IFN-γ) production (5–7). Several inhibitory receptors such as programmed cell death 1 (PD-1), T-cell immunoglobulin domain and mucin domain-3 (TIM3), lymphocyte-activation gene 3 (LAG3), T-cell immune receptor with Ig and ITIM domains (TIGIT), 2B4, and cytotoxic T-lymphocyte associated protein 4 (CTLA4) are also upregulated on their cell surface (3, 4, 8). These changes are typically accompanied by a skewed cellular metabolism. At the early stages of exhaustion, Tex show a limited glucose uptake and a decline in glycolysis, which is ascribed to an elevated expression of PD-1 (9, 10). Defects in mitochondrial mass and fitness are also observed (9, 11). These are attributed to the activation of Blimp-1, which suppresses the expression of PPAR-gamma co-activator 1α (PCG-1α), a key regulator of energy metabolism (11).

Several studies have reported a distinctive transcriptional profile for exhausted CD8 T cells when compared with effector or memory T cells that are differentiated after an acute infection. Indeed, numerous transcription factors have been shown to regulate the maintenance or differentiation of various exhausted CD8 T cell populations. For instance, during chronic infection with Lymphocytic choriomeningitis virus Clone 13 (LCMV Cl13), T-bet negatively regulates PD-1 expression while Eomesodermin (Eomes) is upregulated in antigen specific Tex cells. A high ratio of nuclear Eomes to T-bet positively correlates with the transcription of *Pdcd1* and the level of exhaustion (12). T cell factor-1 (TCF-1) also plays an important role in that it mediates the differentiation of exhausted CD8 T cell precursors (13), negatively regulates the effector-like gene differentiation program, and is upstream of EOMES and c-Myb expression, which regulates cell survival (14). Members of Nuclear factor of activated T cells (NFAT) family of transcription factors also play a center role in T cell exhaustion. NFAT homodimers bind and regulate the expression of inhibitory receptors including *Pdcd1*, *Lag3*, *Havcr2*, thus promoting the Tex phenotype (15, 16). Recent studies have reported an elevated level of thymocyte selection-associated high mobility group box (TOX) expression in exhausted T cells during chronic infection and in tumor-specific T cells (17–21). TOX is a master mediator of T cells exhaustion by governing the expression of inhibitory receptors such as PD-1, TIM3, LAG3, and exhaustion-associated markers such as CD38, and transcription factors including EOMES and TCF-1. TOX expression is induced by persistent TCR activation and NFAT stimulation (18, 19). Other transcription factors, such as Nuclear Receptor Subfamily 4A (NR4A), B lymphocyte-induced maturation protein −1 (Blimp-1), and Basic leucine zipper ATF-like Transcription Factor (BATF) also contribute to Tex development (22–25). Recently, several members of the Interferon regulatory factor (IRF) family were also shown to regulate the CD8 T cell fate. While IRF-4 (26) and IRF-2 (27) appear to be driving CD8 T cell exhaustion, IRF-9 seems to prevent it (28).

Interferon Regulatory Factor 5 (IRF-5) is a transcription factor belonging to the IRF family (29). IRF-5 was originally characterized as a transcription factor downstream of the Type I interferon signaling pathway in innate immune cells (30). Since then, it has been reported to play an essential role in the regulation of many other processes. These include tumor suppression (31), cell cycle regulation (32), induction of apoptosis (33), regulation of proinflammatory cytokines in myeloid lineages (34), M1 macrophage polarization (35), and regulation of plasma cell development (36). Recently, IRF-5 was also shown to be expressed in CD4 T cells during chronic infection and to be downstream of TCR signaling (37). Upon CD4 T cell activation, IRF-5 initiates the TCR assembly and regulates the induction of Th1- and Th17-associated cytokines and the migration of CD4 T cells to the lymph nodes (37). We have recently shown that IRF-5 is also downstream of TLR-7 and induces cell death in murine IFN-γ^+^ CD4 T cells during the chronic visceral leishmaniasis and in memory CD4 T cells of people living with HIV (38, 39). Whether IRF-5 is expressed in CD8 T cells and the role it may have in these cells remains unknown.

Here we show that IRF-5 is upregulated and activated in CD8 T cells in mice infected with LCMV Cl13. In the absence of IRF-5, CD8 T cells enter the cell cycle in greater proportion; however, they fail to accumulate, display reduced survival capacity, and show increased signs of functional exhaustion during the chronic stage of infection, when compared with IRF-5 sufficient cells. Moreover, IRF-5 deficiency in CD8 T cells results in severely defective carbohydrate and lipid metabolism, in a defective mitochondrial envelope, and in the reduced capacity to produce ATP. Finally, we demonstrate that IRF-5-deficiency in CD8 T cells also leads to increased lipid peroxidation and to cell death. Taken together, these results suggest a role for IRF-5 in limiting cell death during chronic LCMV infection by regulating mitochondrial functions and contributing to the metabolic rewiring in CD8 T cells.

## Results

### IRF-5 is mostly expressed and activated in effector and memory-like CD8 T cells

IRF-5 has been shown to be expressed and activated in human and murine CD4 T cells (38, 39); however, whether CD8 T cells also express IRF-5 remains unknown. Hence, we first monitored the expression of this transcription factor in murine CD8 T cells over the course of infection with LCMV Cl13 and found that IRF-5 was indeed expressed by murine CD8 T cells (Figure 1A). As already observed in CD4 T cells (38), this transcription factor was mostly expressed in antigen-experienced cells, while only few cells from naïve mice were IRF-5^+^ (Figure 1B and C). Interestingly, IRF-5 expression peaked around d30p.i. in KLRG1^+^ CD8 T cells, a time when cells start showing signs of exhaustion. At this stage of infection, about 80% of the KLRG1^+^ cells were IRF-5^+^ (Figure 1B). The kinetics of IRF-5 expression in CD127^+^ CD8 T cells was slightly different. Indeed, higher frequencies of IRF-5^+^ CD127^+^ cells were observed between d8 and 34p.i. (Figure 1C). As IRF-5 dimerizes and translocate to the nucleus upon activation (40), we next assessed the subcellular localization of this transcription factor. IRF-5 expression was found to colocalize with the nucleus in CD8 T cells during the whole course of infection, with 60-70% of the cells expressing IRF-5 in the nucleus between d8 and 60 p.i. (Figure 1D). These results suggest that this transcription factor is expressed and is active in effector and memory-like CD8 T cells.

**Figure 1:**
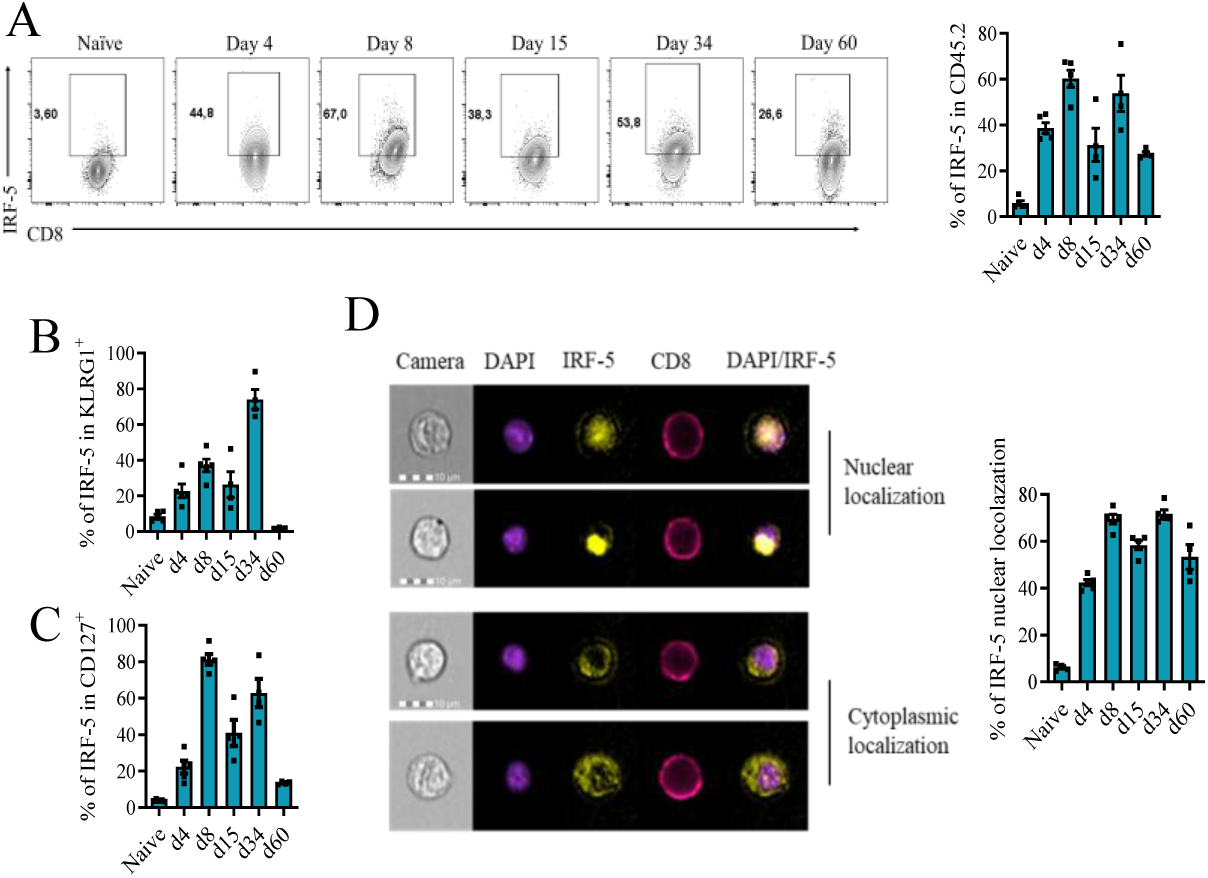
IRF-5 is expressed in both KLRG1^+^ and CD127^+^ CD8 T cells in LCMV Cl13-infected mice. Sorted naive p14 CD8 T cells were transferred into CD45.1 recipient mice one day prior to intravenous infection with 2 × 10^6^ PFU LCMV Cl13. Mice were euthanized at various time points after infection. Graphs show (A) representative flow cytometry plots (left) and the frequency (right) of p14 CD8 T cells present in the spleen of infected mice; (B) the percentage of KLRG1^+^ CD127^-^ p14 CD8 T cells expressing IRF-5; (C) the percentage of CD44^+^ CD127^+^ p14 CD8 T cells expressing IRF-5; and (D) the frequency of IRF-5 expression that colocalizes with DAPI staining over the course of LCMV Cl13 infection. Data represent the mean ± SEM of one of 2 or 3 independent experiments, n = 4-5.

### IRF-5 is required to maintain CD8 T cells during chronic infection

To determine the function of IRF-5 in CD8 T cells, we generated IRF-5-deficient p14 transgenic mice (*Irf5^flox/flox^* x *CMV-Cre^+^* p14), which have CD8 T cells that are specific for the LCMV gp33 antigen. We first confirmed IRF-5 deletion in total splenocytes (Supplemental Figure 1A) and in splenic CD8 T cells (Supplemental Figure 1B) from *Irf5^-/-^* p14 mice by flow cytometry. Naïve CD8 T cells purified from these mice and from the IRF-5 sufficient, *Cre^-^* p14 control mice (*Irf5^flox/flox^*x *CMV-Cre^-^ p14*; herein called as WT p14) were then adoptively transferred into CD45.1 congenic mice a day prior to infection with LCMV Cl13 (Supplemental Figure 1C). Adoptive transfer experiments were carried out using separate groups of male and female mice. We found that male and female mice that received WT p14 cells had different survival rates (Supplemental Figure 1D), with only 60% of male mice surviving LCMV Cl13 infection compared to 80% of the females. However, the most remarkable difference was observed between male and female mice that received *Irf5^-/-^* p14 cells. While females adoptively transferred with *Irf5^-/-^* p14 cells survived infection to similar rate than those that received WT p14 cells, only 20% of male mice adoptively transferred with *Irf5^-/-^* p14 survived LCMV Cl13 infection at d15 p.i. (Supplemental Figure 1D). Because of these differences, we decided to characterize the function of IRF-5 in CD8 T cells from female and male mice separately. To this end, p14 cells were adoptively transferred into recipient mice and monitored at various time points over 60 days of infection. As expected, WT p14 cell expansion peaked at d8 p.i. to slightly contract during the chronic stage of infection (Figure 2A). This kinetics was reflected in the percentages of p14 cells (Figure 2B) as well as in the p14 cell numbers (Figure 2C) observed over the course of infection. Surprisingly, IRF-5-deficient p14 cells underwent clonal expansion similarly to WT p14 cells but failed to survive during the chronic stage of infection (Figure 2A). Indeed, a dramatic decrease in cell frequencies (Figure 2B) and numbers (Figures 2C) was observed at d30 and 60 p.i. in mice that received IRF-5-deficient p14 cells compared to the control group. Interestingly, IRF-5 deficiency mostly impacted KLRG1^+^ cells, which were present at significantly lower frequencies in the spleen of mice that received *Irf5^-/-^* p14 compared to WT p14 cells at d30 and 60 p.i. (Figure 2D and E). In contrast, noticeably higher frequencies of KLRG1^+^ cells were observed at d8 p.i. in the spleen following adoptive transfer of *Irf5^-/-^* p14 compared to WT p14 cells (Figure 2D and E). CD127^+^ memory-like cells were less affected by the lack of IRF-5 (Figure 2F). We found similar results in female mice, where IRF-5-deficiency resulted in higher percentages of KLRG1^+^ cells at d8 p.i. compared with WT p14 cells but KLRG1^+^ responses were not maintained to the same level in the absence of IRF-5 (Supplemental Figure 1 E). Taken together, these results suggest that IRF-5 deficiency in p14 cells results in *Irf5^-/-^*p14 cells undergoing similar clonal expansion than WT p14 cells but then failing to survive during the chronic phase of infection.

**Figure 2:**
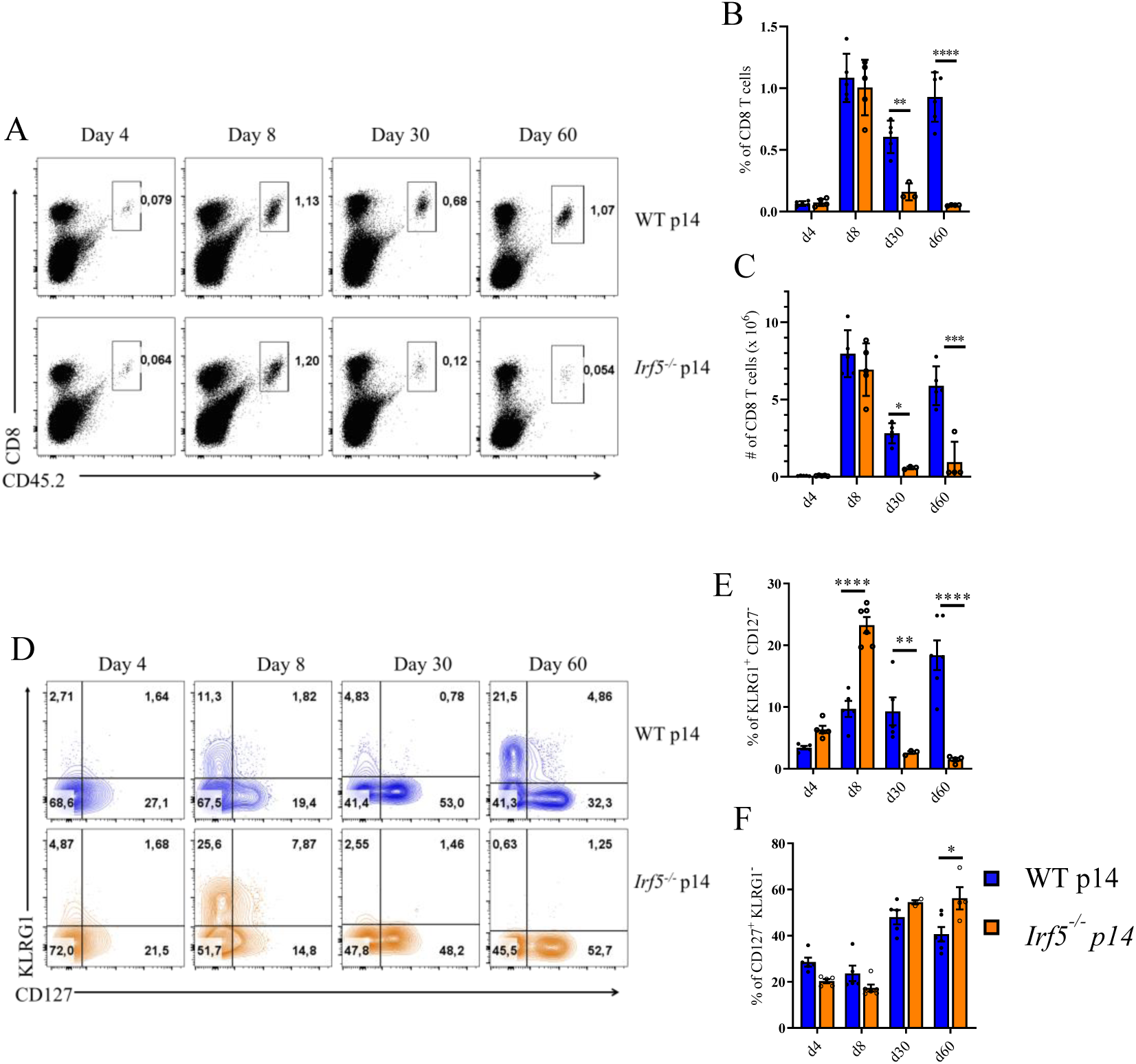
IRF-5 is required to maintain CD8 T cell responses throughout infection. CD8 T cells from male WT or *Irf5^-/-^* p14 mice were transferred into CD45.1 mice one day prior to intravenous infection with 2 × 10^6^ PFU LCMV Cl13. Mice were euthanized at various time points after infection. Graphs show (A) representative FACS plots for splenocytes stained with anti CD8 and anti CD45.2; the percentage (B) and the absolute numbers (C) of adoptively transferred WT and *Irf5^-/-^* p14 CD8 T cells present in the spleen of infected recipient mice; (D) representative flow cytometry plots and corresponding percentages of KLRG1^+^ CD127^-^ (E) and CD127^+^ CD44^+^ (F) p14 CD8 T cells in the spleen of recipient mice over the course of LCMV infection. Data represents the mean ± SEM, n=4-5, one of 3 independent experiments is shown. * denotes *p*<0.05, ** denotes *p*<0.01 *** denotes p< 0.001 **** denotes *p*< 0.0001.

To confirm our results in non-transgenic CD8 T cells, we infected *Irf5^flox/flox^* x *Lck-Cre*^+^ and *Cre*^-^ mice (herein referred to as *Lck-Cre*^+^ and WT) with LCMV Cl13 and monitored CD8 T cell responses over the course of infection. A more pronounced decline in percentages (Supplemental Figure 2A and B) and numbers (Supplemental Figure 2C) of total CD8 T cells was observed from d28 p.i. in mice with a T cell-specific IRF-5 ablation compared with the control group, indicating that IRF-5 deficiency affected endogenous, non-transgenic cell survival as well.

### *Irf5* ablation results in functional exhaustion of CD8 T cells

We next investigated whether the absence of IRF-5 in p14 cells affected their effector function. Therefore, the production of various cytokines was assessed at different time points of infection upon restimulation. We found that *Irf5^-/-^* p14 cells have a superior capacity to generate IFN-γ-producing cells at d8p.i. in terms of frequencies (Figure 3 B, upper panel) and cytokine amount (Figure 3B, lower panel), compared with WT p14 cells. Similar results were observed for TNF-(Figure 3C) and INF-γ-TNF-double producing cells (Figure 3D), reflecting our previous observation that mice that received *Irf5^-/-^* p14 displayed higher frequencies of KLRG1^+^ effectors at d8 p.i. (Figure 2E). Nevertheless, the absence of IRF-5 significantly impacted the maintenance of IFN-γ-producing cells over the chronic stage of infection, when lower frequencies of IFN-γ^+^ cells (Figure 3A and 3B, upper panel) and lower amounts of IFN-γ per cell (Figure 3B, lower panel) were observed, compared with the control group. The percentage of TNF^+^ (Figure 3A and 3C) and IFN-γ ^+^TNF^+^ (Figure 3A and 3D) *Irf5^-/-^*p14 cells was also significantly lower than the control group at d30 and d60 p.i.. Interestingly, mice that received *Irf5^-/-^* p14 cells showed reduced frequencies of IL-2-producing cells over the whole course of infection compared with those that were adoptively transferred with WT p14 (Figure 3A and 3E, upper panel). Moreover, *Irf5^-/-^* p14 cells produced less IL-2 than WT p14 cells (lower panel) and dramatically lower percentages of IFN-γ^+^ TNF^+^ IL-2^+^ were observed for *Irf5^-/-^* p14 compared with WT p14 cells (Figure 3A and 3F). Similar results were obtained during the chronic phase of infection using female mice (Supplemental Figures 3A-C).

**Figure 3:**
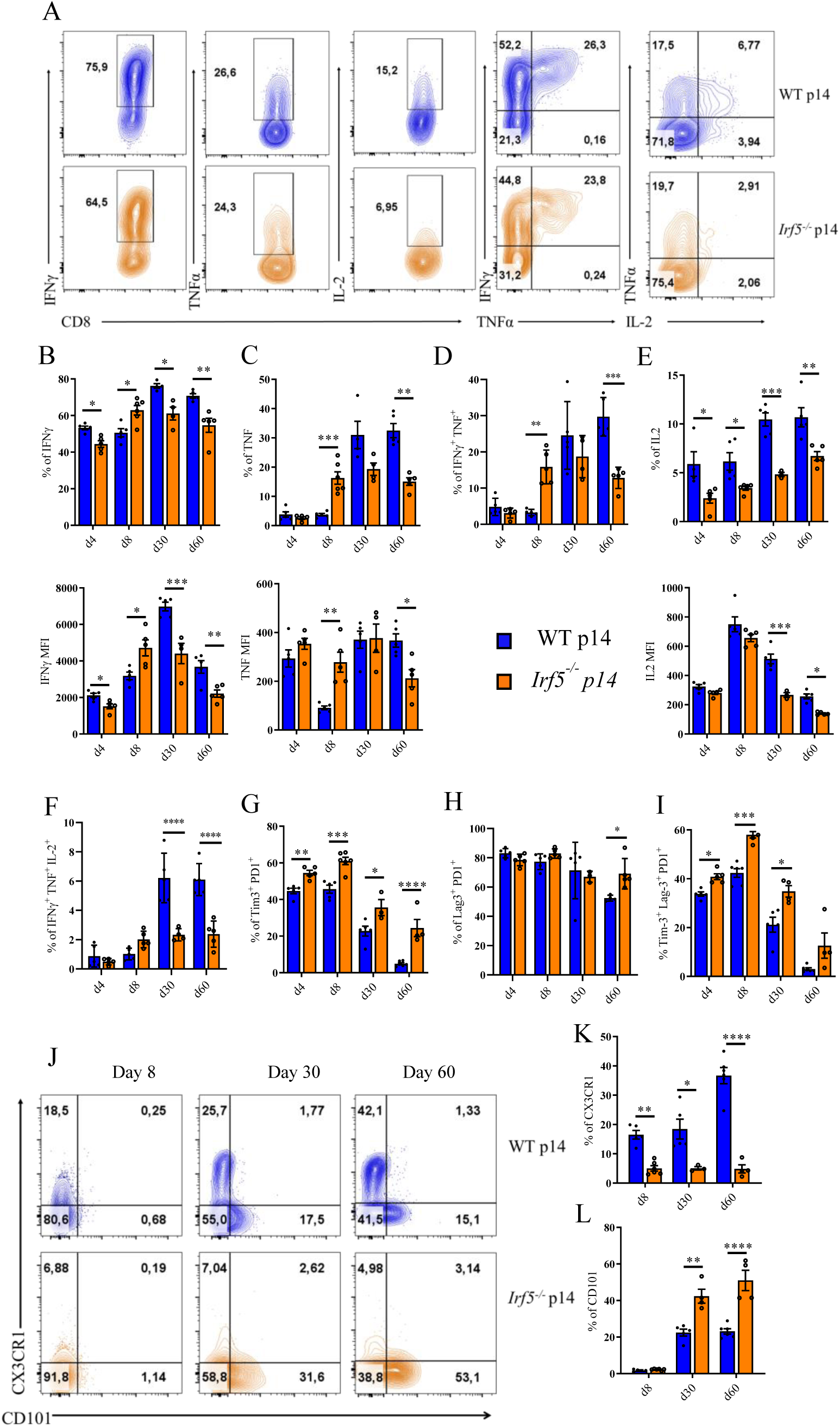
IRF5 ablation is associated with functional exhaustion during chronic infection. CD8 T cells from male WT or *Irf5^-/-^* p14 mice were adoptively transferred into CD45.1 recipient mice one day prior to intravenous infection with 2 × 10^6^ PFU LCMV Cl13. Mice were euthanized at various time points after infection. (A-F) Cytokine production by adoptively transferred p14 cells was assessed after restimulation with the gp33 peptide. (A) Representative FACS plot for the various cytokine staining at d30 p.i.. (B-D) Graphs show the frequency (upper graphs) and mean fluorescence intensity (MFI) (lower graphs) of IFN-γ^+^ (B), TNF^+^ (C), and IL-2^+^ (D) WT p14 and *Irf5*^-/−^ p14 CD8 T cells at various days post infection. (E-F) Graphs represent the percentage of IFN-γ^+^ TNF^+^ (E) and IFN-γ^+^ TNF^+^ IL-2^+^ (F) WT p14 and *Irf5*^-/−^ p14 CD8 T cells at various days post infection. (G-I) Expression of inhibitory receptors in WT and *Irf5*^-/−^ p14 CD8 T cells. Graphs represent the frequency of Tim3^+^ PD1^+^ (G), Lag3^+^ PD1^+^ (H), and Tim3^+^ Lag3^+^ PD1^+^ (I) adoptively transferred WT p14 and *Irf5*^-/−^ p14 cells over the course of infection. (J-L) Representative flow cytometry plots and graphs showing the percentage of CX3CR1^+^ effector-like (J and K) and CD101^+^ terminally exhausted (J and L) WT and *Irf5*^-/−^ p14 CD8 T cells at various time points post infection. Data represent the mean ± SEM, n=4-5, one of 3 independent experiments is shown, * denotes *p*<0.05, ** denotes *p*<0.01 *** denotes p< 0.001 **** denotes *p*< 0.0001.

IRF-5 was shown to compete with IRF-4 (41), which in turn is known to drive CD8 T cell exhaustion (26). Thus, we next assessed whether the absence of IRF-5 expression affected at all IRF-4 expression. Interestingly, we did not observe any differences in the frequency of CD8 T cells expressing IRF-4 between the two groups at any time points observed (Supplemental Figure 4A). Likewise, the proportion of CD8 T cell expressing Eomes (Supplemental Figure 4B) and T-bet (Supplemental Figure 4C) was not altered in the absence of IRF-5, with the exception of d30 p.i., when *Irf5^-/-^*p14 cells expressed significantly lower levels of T-bet compared to their WT counterparts (Supplemental Figure 4C).

Because these findings suggest that *Irf5^-/-^* p14 cells are more dysfunctional at d30 and 60 of infection than their IRF-5-sufficient counterpart, we next assessed whether these cells would show increased signs of exhaustion. First, we assessed the expression of three inhibitory receptors that are typically expressed by exhausted CD8 T cells, namely PD-1, TIM-3, and LAG-3 (42). We observed significantly higher frequencies of cells co-expressing PD-1 and TIM-3 in IRF-5-deficient p14 cells over the entire course of infection, compared with the control group (Figure 3G). Percentages of PD-1^+^ LAG-3^+^ cells were very similar in both study groups with exception of d60p.i., when these cells’ frequency was higher in the *Irf5^-/-^* p14 group compared with the control group (Figure 3H). However, triple positive cells (PD-1^+^ Tim-3^+^ Lag-3^+^) were significantly more present in the absence of IRF-5 (Figure 3I), suggesting that IRF-5 deficiency could worsen CD8 T cell exhaustion. To confirm this possibility, we assessed cell surface modulation of the chemokine receptor CX3CR1 and the transmembrane glycoprotein CD101. These two markers allow to differentiate exhausted (CD101^+^) from effector-like (CX3CR1^+^) CD8 T cells during chronic infections (43). As expected, we found dramatically reduced frequencies of CX3CR1^+^ effector-like cells (Figure 3J and K) and a significantly higher percentage of CD101^+^ exhausted cells (Figure 3J and L) in the absence of IRF-5 when compared with the IRF-5 sufficient group. Similar results were obtained using female mice (Supplemental Figures 3D-E). We also analyzed effector function and inhibitory receptor expression in T cell-specific *Irf5^-/-^* mice infected with LCMV Cl13 and found that in the absence of IRF-5, stronger signs of functional exhaustion (Supplemental Figures 5A-C) and higher frequencies of inhibitory receptors’ expressions (Supplemental Figures 5D-F) were observed in CD8 T cells, compared with the control group. Interestingly, mice with a T cell specific *Irf5* ablation failed to clear infection in the serum (Supplemental Figures 5G), highlighting the severe CD8 T cell dysfunctionality.

### IRF-5 deficiency alters the expression of genes involved in the cell cycle

To further characterize the role of IRF-5 in CD8 T cells, we compared the transcriptomic profile of IRF-5-sufficient and -deficient p14 cells by bulk RNA sequencing. To this end, we performed adoptive transfer experiments as described in Supplemental Figure1C, using male and female mice. Adoptively transferred cells were purified from the spleen of mice infected with LCMV Cl13 at d21 p.i. and bulk RNA sequencing was performed on two groups of male and female mice, representing a pool of 7-17 mice each. The top up- and down-regulated genes are shown in Supplemental Figure 6 and Supplemental Table 1. We first performed gene ontology analysis to identify the biological processes that were mostly dysregulate. We found that IRF-5 deficiency predominantly affected the expression of genes involved in the mitotic cell cycle and cell division process (Figure 4A), particularly those involved in accelerating cell division (Figure 4B). These included *Mki67*, a marker for cell proliferation, and other cell cycle-associated genes such as *Cdca5*, *Cdca8*, and *Cdc45* (Figure 4B). These results were not surprising, given the known role of IRF-5 in cell cycle arrest (32). Hence, we decided to validate these findings *in vivo* and analyze the proportion of adoptively transferred IRF-5-sufficient and -deficient p14 cells that were in the G0, G1, and S-G2-M phase at d4, 8, 30, and 60 after infection with LCMV Cl13. We observed a significantly higher proportion of *Irf5^-/-^* p14 cells that were in the G1 and S-G2-M phase at d30 and 60 p.i., when compared with WT p14 cells (Figure 4C). This suggests that *Irf5^-/-^* p14 cells were indeed entering the cell cycle in greater proportion. To monitor whether this would also occur *in vitro*, we stimulated purified naïve *Irf5^-/-^* and WT p14 cells *in vitro* with the gp33 peptide and monitored the proportion of cells in the G0, G1 and S-G2-M phase 6, and 24h after stimulation. No differences were observed between both groups in the absence of stimulation (Figure 4D). As expected, *Irf5^-/-^* p14 cells were entering the cell cycle in greater proportion than their WT counterpart already at 6h after stimulation with the gp33 peptide *in vitro* as well; this difference was sustained over the first 24h of culture (Figure 4D). This suggests that the absence of IRF-5 leads to enhanced cell division. Nevertheless, even though they enter the cell cycle more rapidly, the proportion and numbers of *Irf5^-/-^* p14 cells found in the spleen of LCMV Cl13-infected mice at d30 and 60p.i. is significantly lower than WT p14 cells (Figure 2B and C).

**Figure 4:**
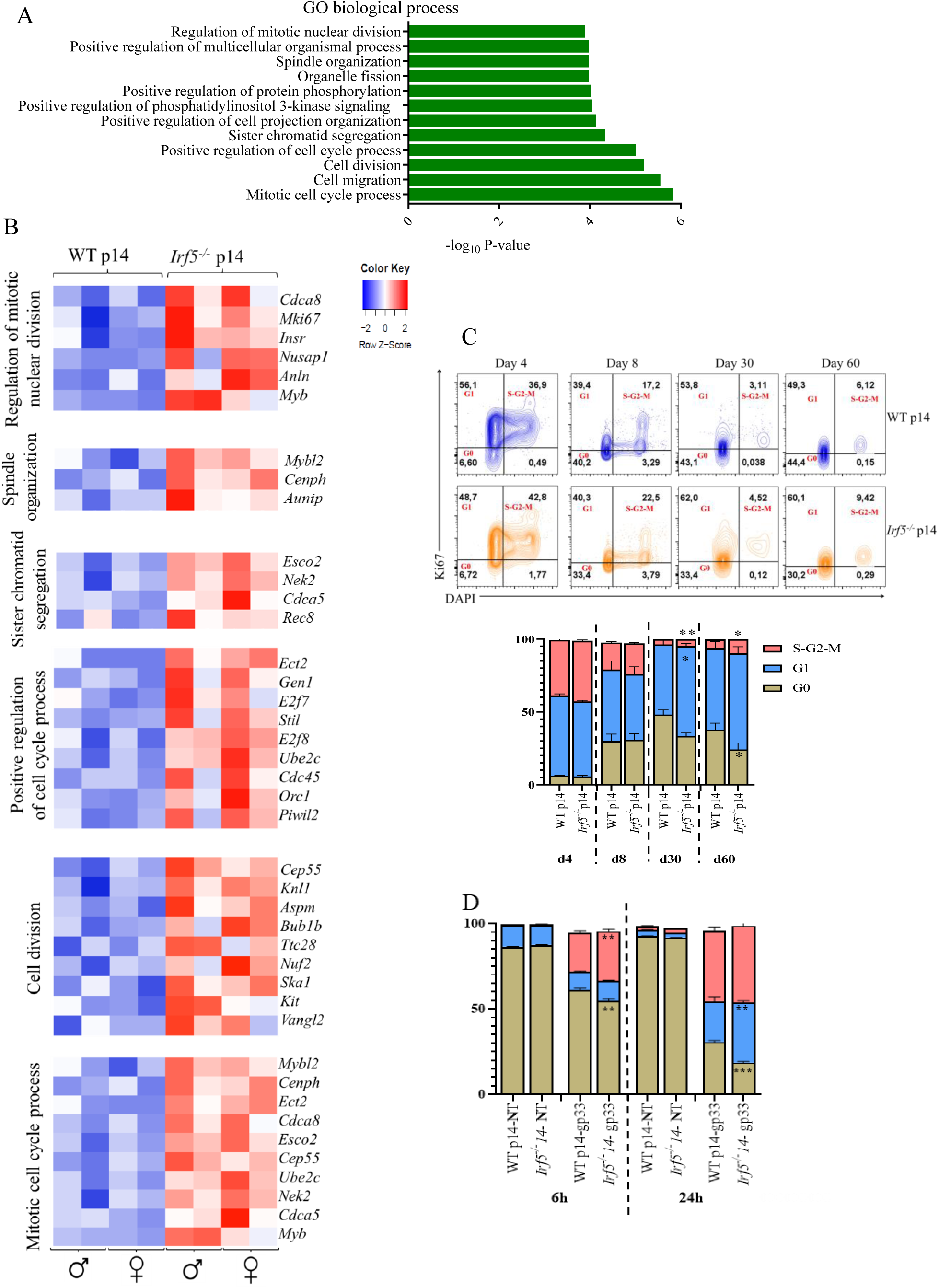
IRF-5-deficiency alters the expression of genes involved the cell cycle. (A-B) Bulk RNA sequencing was performed on adoptively transferred WT p14 and *Irf5^-/-^* p14 CD8 T cells isolated from male and female recipient mice at day 21 after LCMV Cl13 infection. (A) Gene ontology analysis results showing biological processes that were mostly upregulated in the absence of IRF-5. (B) Heat map illustrating the relative expression of genes involved in the cell cycle in WT and *Irf5^-/-^* p14 CD8 T cells. (C) WT and *Irf5^-/-^* p14 CD8 T cells were adoptively transferred into CD45.1 recipient mice a day prior to infection with LCMV Cl13. Mice were then euthanized at various time point after infection. Graphs show representative flow cytometry plots for Ki67 and DAPI staining (upper graphs) and the proportion of cells in the various phases of the cell cycle at any given time point (lower graph). (D) Graph displays the cell cycle analysis based on Ki67 and DAPI staining of WT and *Irf5^-/-^* p14 CD8 T cells stimulated *in vitro* with or without the gp33 peptide. Data represent the mean ± SEM, n=4-5, one of 3 independent experiments is shown, * denotes *p*<0.05, ** denotes *p*<0.01.

### IRF-5 deficiency in CD8 T cells affects the cellular metabolism

To better understand why *Irf5^-/-^* p14 cells fail to accumulate, we went back to our RNA sequencing data and performed Gene Ontology analysis of various pathways that could compromise cell survival or cell proliferation. First, we reasoned that *Irf5^-/-^* p14 cells were probably dying more significantly than their WT counterpart. Indeed, several genes associated with cell death (GO:0008219) were more significantly upregulated in p14 cells in the absence of IRF-5, particularly in cells from female mice, suggesting that cell death may occur more frequently in *Irf5^-/-^* than in WT p14 cells at d21 p.i. (Figure 5A). To better understand the cause of cell death, we next assessed whether cell exhaustion pathways (as described by Wherry *et al*., (42)) were enhanced in *Irf5^-/-^* compared with WT p14 cells. We found that, although *Irf5^-/-^* p14 cells showed heightened signs of functional exhaustion *in vivo*, only some of the genes typically associated with exhaustion in CD8 T cells were clearly upregulated compared with WT p14 cells but others were not (Figure 5B), suggesting that functional exhaustion and cell death may results from some other pathway(s).

**Figure 5:**
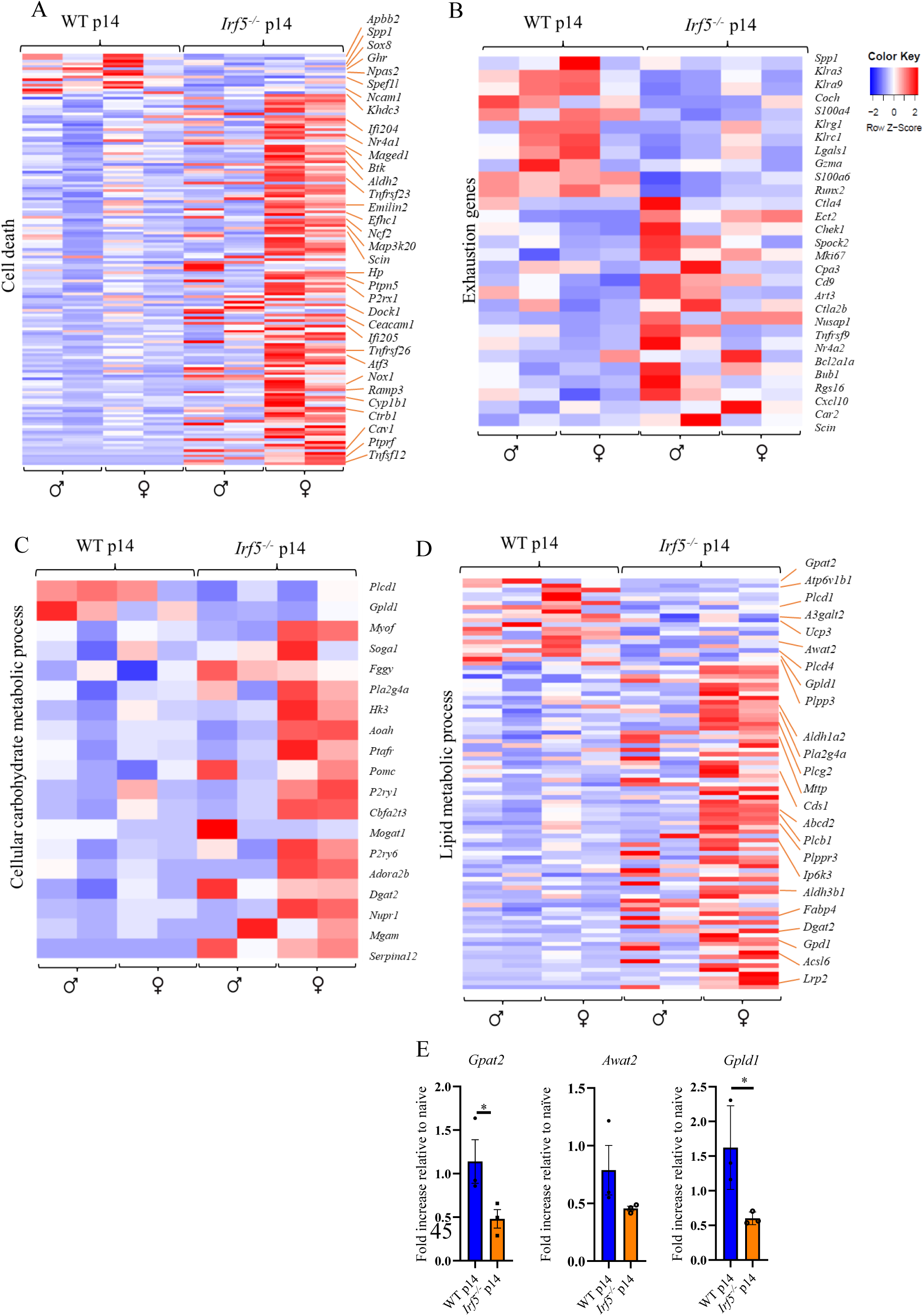
IRF-5-deficiency affects the expression of genes involved in the cellular metabolism. Transcriptional profiling was performed on adoptively transferred WT p14 and *Irf5^-/-^* p14 CD8 T cells isolated from male and female recipient mice at day 21 post LCMV Cl13 infection. (A-D) Heatmaps displaying expression of relative upregulated and down regulated signature gene sets involved in (A) CD8 T cell exhaustion, (B) cell death, (C) cellular carbohydrate metabolic processes, and (D) lipid metabolic processes. (E) WT and *Irf5^-/-^* p14 CD8 T cells were adoptively transferred into female CD45.1 recipient mice a day prior to infection with LCMV Cl13. Adoptively transferred cells were then sorted at d21 p.i and the mRNA expressions of *Gpat2* (left graph), *Awat2* (center graph), and *Gpld1* (right graph) were analyzed by digital droplet PCR. Data represents the mean ± SEM, * denotes *p*<0.05.

Hence, we assessed possible enrichments in gene signatures associated with cellular metabolism. Because IRF-5 was shown to indirectly regulate glycolysis in macrophages upon TLR stimulation (44, 45), we first analyzed genes associated with the carbohydrate metabolism. We found an enrichment of genes associated with the cellular carbohydrate metabolic process (GO: 0044262) in IRF-5-deficient p14 cells compared with their WT counterpart, particularly in cells from female mice (Figure 5C). Some genes were directly associated to glucose metabolism, for example *Hk3*, which encodes for a hexokinase 3, an enzyme that phosphorylates glucose to trap it inside the cell and prepare it for the glycolytic pathway (46); of *Fggy*, which encodes for proteins that phosphorylate carbohydrates for energy production and biosynthetic pathways (47); or the maltase-glucoamylase encoding gene *Mgam* (Figure 5C). Another gene, *P2ry1*(encoding for G-protein-coupled receptor) mostly regulates cell signaling and calcium mobilization (Figure 5C). However, heightened expression of these genes could simply be a consequence of IRF-5-deficient p14 cells entering more rapidly the cell cycle and requiring more energy but does not necessarily explain why cells are dying. Thus, we next assessed the lipid metabolism (GO:0006629). Interestingly, we found several genes that were expressed to a lower degree in *Irf5^-/-^* compared to WT p14 cells. For example, *Gpat2*, which encodes for glycerol-3-phosphate acyltransferase 2, an enzyme involved in triacylglycerol (TAG) synthesis (48); *Awat2*, which encodes for acyl-CoA wax alcohol acyltransferase 2, an enzyme that catalyzes the synthesis of wax esters (49); *Atp6v1b1*, which encodes for a vacuolar ATPase that has been shown to regulate oxidative phosphorylation, and sphingolipid, fatty acid, and energy metabolism, and to severely affect mitochondrial respiration in zebra fish (50); *Ucp3*, which encodes for the uncoupling protein 3, a mitochondrial fatty acid anion exporter that protects mitochondria from lipid-induced damage (51); and *Gpdl1*, which encodes for glycosylphosphatidylinositol-specific phospholipase D, an enzyme involved in triglyceride metabolism (Figure 5D). We confirmed the expression of some of the genes in WT and *Irf5^-/-^* p14 cells purified *ex-vivo* from the spleen of female mice at d21 p.i. and found that *Irf5^-/-^* p14 cells had a significantly lower expression of *Gpat2*, *Awat2*, and *Gpdl1* mRNA levels when compared with the control group (Figure 5E). Together, the transcriptome analysis of IRF-5-deficient p14 cells revealed that these cells undergo cell death during the chronic stages of infection possibly because a defect in their metabolism.

### IRF-5 deficiency profoundly affects cell respiration and ATP production

We next analyzed the bioenergetic flux in live cells to investigate the oxidative metabolism as well as the glycolysis of female WT and *Irf5^-/-^* p14 cells stimulated or not with the gp33 peptide. Thus, we quantified the oxygen consumption rate (OCR) (Figure 6A) and the extracellular acidification rate (ECAR) (Figure 6B) of both groups of cells at 6 and 24 hours after stimulation with the gp33 peptide. We found profound defects in the respiratory capacity of IRF-5-deficient p14 cells, which had a severely compromised basal (Figure 6A and C) and maximal (Figure 6A and D) respiration. This was particularly noticeable 24h after stimulation. The ATP production derived from OCR was also dramatically reduced compared with WT p14 cells (Figure 6E). When we analyzed the bioenergetic profile at 24h after stimulation, we found that WT p14 cells stimulated with the gp33 peptide could gain ATP from both mitochondrial respiration and glycolysis, whereas IRF-5-deficient p14 cells were nearly unable to produce ATP through both pathways (Figure 6F). This was not due to their incapacity to uptake glucose, because *Irf5^-/-^* p14 cells could internalize the fluorescent glucose analog 2-NBDG at levels similar to the IRF-5 sufficient cells (Figure 6G). Moreover, IRF-5 deficiency did not impair the capacity to uptake lipids either. Indeed, *Irf5^-/-^* p14 cells were superior in internalizing fatty acids 6h and 24h after stimulation with the gp33 peptide, when compared with their WT counterpart (Figure 6H). Similar results, although less dramatic, were obtained using male WT and *Irf5^-/-^* p14 cells (Supplemental Figures 7A-H). In sum, our results indicate that IRF-5-deficiency severely impairs CD8 T cell respiration and energy production.

**Figure 6:**
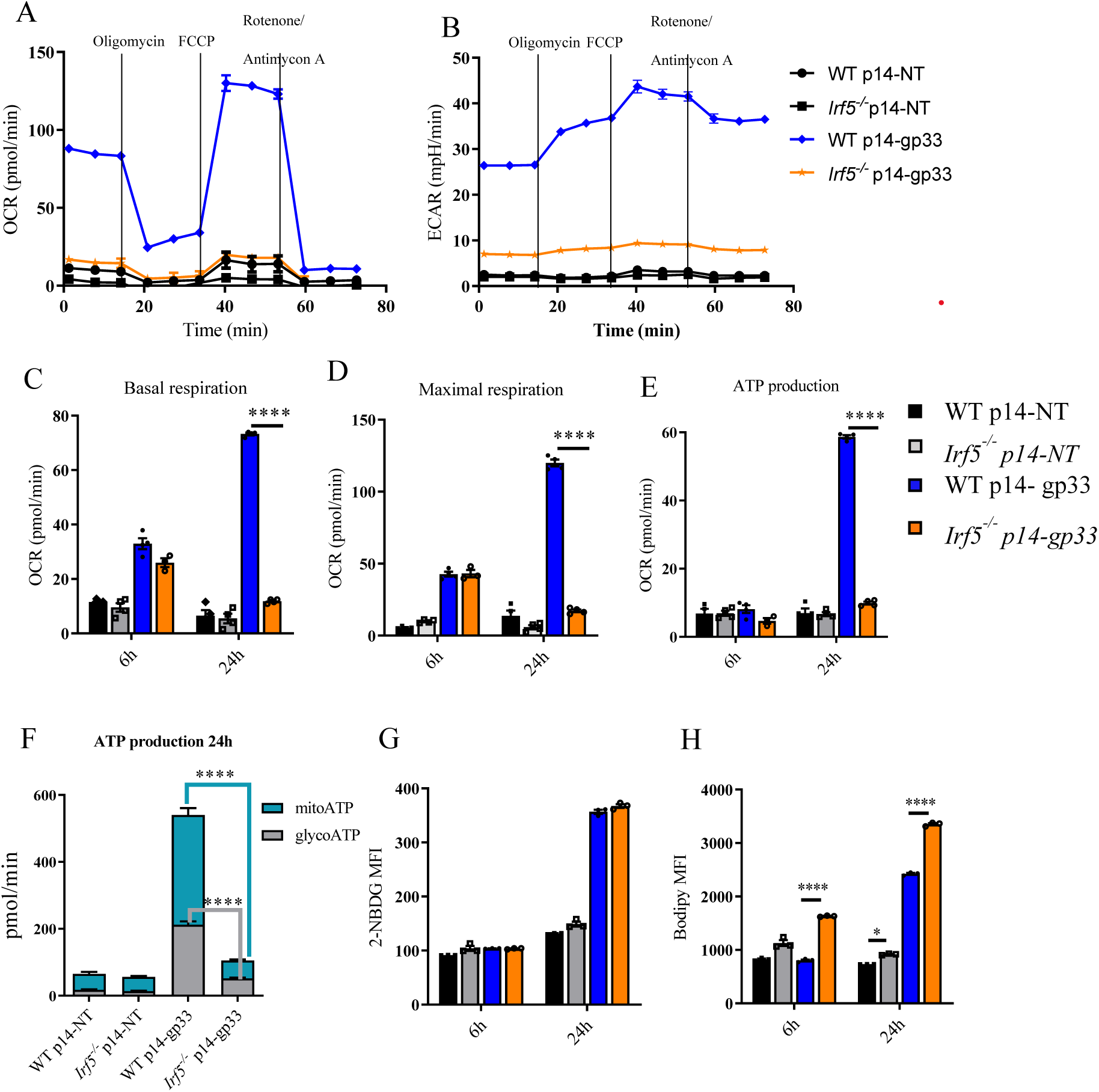
IRF-5-deficient p14 cells show profound defects in the cellular respiration and ATP production. WT p14 and *Irf5^-/-^* p14 CD8 purified from female mice were cultured *in vitro* in the presence or absence of the gp33 peptide, and the mitochondria respiratory capacity was measured using the Seahorse XFe-96 analyzer. Graphs illustrate (A) the representative oxygen consumption rates (OCR) and (B) the extracellular acidification rates (ECAR) over time; (C) the basal and (D) maximal respiration, and (E) the ATP production at 6h and 24h after stimulation with the gp33 peptide; (F) the ATP production rate; and (G) the glucose (measured using the fluorescent glucose analog 2-NBDG) and (H) the fatty acid uptake capacity (quantified using bodipy FLC_16_ fluorescence intensity). Data represent the mean ± SEM, n=4, one of three independent experiments is shown, * denotes *p*<0.05, **** denotes *p*<0.0001

### IRF-5 regulates the mitochondrial electron transport chain

To test whether the dysregulated cellular metabolism observed in IRF-5-deficient cells was a consequence of an intrinsic mitochondrial defect, we analyzed the expression of genes associated with the mitochondria envelope (GO:0005740). Interestingly, we found several differentially expressed genes between the two groups of study (Figure 7A). Several genes were expressed at lower levels in *Irf5^-/-^* compared with WT p14 cells, among those we found: *Gpat2*, *Ucp3*, *Coq4*, *Ndufaf3*, and *Ndufb4* (Figure 7A). *Coq4* encodes for the coenzyme Q4, a protein involved in the biosynthesis of an essential component of the electron transport chain in mitochondria (52); *Ndufaf3* encodes for the NADH hydrogenase 1 alpha subcomplex assembly factor, a protein involved in the assembly and function of the mitochondria respiratory chain complex I (53); *Ndufb4* encodes for the NADH hydrogenase 1 beta subcomplex 4, a subunit of the mitochondrial respiratory chain complex I (54). We validated the results by assessing the mRNA expression levels of *Coq4*, *Ndufaf3*, and *Ndufb4* in WT and *Irf5^-/-^*p14 cells purified *ex-vivo* from female LCMV Cl13-infected mice at d21 p.i.. We found that indeed IRF-5-deficient p14 cells expressed significantly lower levels of those three genes when compared with IRF-5-suffiecient p14 cells (Figure 7B). Among the upregulated genes, we could not find the gene encoding for the Growth hormone inducible transmembrane protein (GHITM) (Figure 7A). A recent study by Orliaguet *et al*. has shown that IRF-5 regulates the mitochondrial architecture remodelling by transcriptionally repressing a key mitochondrial component for oxidative respiration, namely GHITM (55). This work was done in macrophages and in the context of a high-fat diet. However, when we analyzed *Ghitm* mRNA levels in *ex-vivo* purified cells, we observed a significant upregulation of this gene’s expression in IRF-5-deficient compared with IRF-5-sufficient cells (Figure 7C).

**Figure 7:**
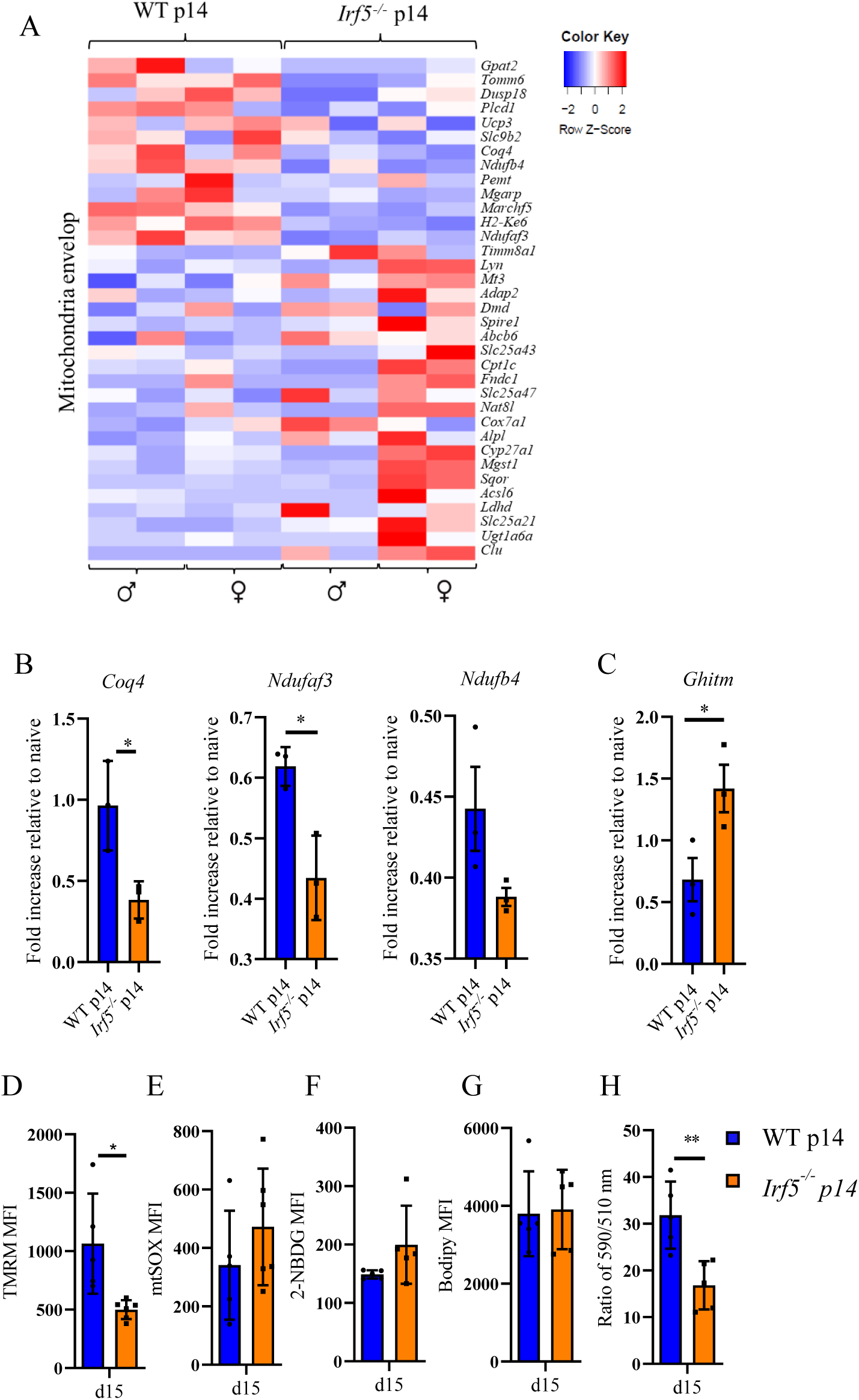
IRF-5-decificiency mostly affects the mitochondrial electron transport chain and leads to lipid peroxidation. (A) Bulk RNA sequencing data of adoptively transferred WT and *Irf5^-/-^* p14 CD8 T cells isolated from recipient female and male mice at d21 p.i. were analyzed for the expression of signature gene sets associated with the mitochondrial envelop. The heatmap illustrate up- and down-regulated genes. (B-H) WT or *Irf5^-/-^* p14 CD8 T cells were adoptively transferred into female CD45.1 mice a day prior to infection with LCMV Cl13. Mice were then euthanized at d21 p.i (B-C) and d15 p.i. (D-H) and adoptively transferred cells were sorted. Graphs represent (B) mRNA expressions of *Coq4* (left graph), *Ndufaf3* (center graph), *Ndufb4* (right graph) and (C) *Ghitm* measured by digital droplet PCR; (D) the mitochondrial membrane potential, determined by TMRM mean fluorescence intensity; (E) the mitochondrial ROS production, assessed by mtSOX fluorescence intensity; (F) the glucose uptake capacity, measured using the glucose analog 2-NBDG; (G) the bodipy FLC_16_ mean fluorescence intensity as an indicator of the fatty acid uptake capacity; and (H) the relative cellular lipid peroxidation level, detected using Image-iT™ Lipid Peroxidation Kit and calculated using the ratio of the signal from the 590 nm and 510 nm channels (lower ratios indicate higher lipid peroxidation levels). Data represent the mean ± SEM, n=5, * denotes *p*<0.05, ** denotes *p*<0.01

So far, our results indicate that *Irf5^-/-^* p14 cells have a severe defect in the mitochondrial envelope, which impairs mitochondrial functions. To confirm that mitochondria were not functioning optimally in the absence of IRF-5, we next measured the mitochondrial membrane potential *ex-vivo* in cells from LCMV Cl13 -infected female mice adoptively transferred with WT and *Irf5^-/-^* p14 cells. Indeed, we noticed a fluorescence drop in *Irf5^-/-^* compared with WT p14 cells (Figure 7D), suggesting that these cells were metabolically stressed. Moreover, we observed a noticeable, yet not significant difference in mitochondrial reactive oxygen species (ROS) levels *ex-vivo* in *Irf5^-/-^* compared with WT p14 cells (Figure 7E). As previously observed in *in vitro* stimulated cells (Figure 6G), IRF-5 deficiency did not alter the cells’ capacity to internalize glucose (Figure 7F) or fatty acids (Figure 7G). Hence, we reasoned that if *Irf5^-/-^* p14 cells can internalize lipids equally as well as WT p14 cells and ROS levels are elevated, this means that perhaps lipid peroxidation occurs in those cells. To this end, we measured lipid peroxidation *ex-vivo* and found, indeed, that lipid peroxidation was occurring in IRF-5-deficient p14 cells at d21 p.i. (Figure 7H). In summary, our results demonstrate that in the absence of IRF-5, CD8 T cells enter the cell cycle more rapidly, but are metabolically stressed because of a severe defect in the mitochondrial envelope and die possibly by lipid peroxidation.

## Discussion

In the past few years, several studies have uncovered numerous transcription factors governing the intricate gene expression program underlying CD8 T cell exhaustion during chronic infections. However, relatively little is known about factors that help CD8 T cell to survive in the hostile chronic inflammatory environment typical for persistent infections. Here, we show that IRF-5 prevents premature functional exhaustion and cell death by controlling the cell cycle and contributing to the metabolic rewiring of CD8 T cells during chronic infection. Indeed, IRF-5-deficient CD8 T cells quickly become functionally exhausted, display severe defects in mitochondrial functions and lipid metabolism, and ultimately die, possibly as a consequence of lipid peroxidation.

IRF-5 function has been studied in detail in antigen presenting cells and tumor cells, where it regulates the inflammatory and anti-viral response (56), macrophage polarization (35), cell cycle (32), and cell death (33, 57). Recently, a function for IRF-5 in governing the carbohydrate metabolism was also ascribed (44, 45). In contrast, the literature on the role of IRF-5 in T cells is sparse and only describes IRF-5 functions in CD4 T cells (37, 38). We have now shown that IRF-5’s role substantially differs between CD4 and CD8 T cells. Other transcription factors also have opposing functions in both T cell subsets. For instance, TOX plays an essential role in the CD4 T cell development (58), but drives exhaustion in CD8 T cells (17–21). In our infection model, the absence of IRF-5 profoundly affected CD8 T cell responses to an extent that they are incapable of clearing LCMV Cl13 infection in the serum. Unsurprisingly, we found a dysregulated cell cycle in IRF-5-deficient CD8 T cells. Indeed, IRF-5 was shown to exert cell cycle arrest functions in a p53-dependent (59) and -independent (33) way in tumor cells. In LCMV Cl13-infected mice, CD8 T cells were entering the cell cycle more rapidly in the absence of IRF-5; however, IRF-5 ablation did not affect the clonal expansion, as similar numbers and frequencies of CD8 T cells were observed at d8 p.i. in WT and IRF-5 deficient cells. Moreover, *Irf5^-/-^* p14 CD8 T cells were dying more rapidly than their WT counterparts.

In the search for reasons for a premature cell death, we found that *Irf5^-/-^* p14 CD8 T cells were not capable to cope metabolically and gain energy through the carbohydrate and the lipid metabolism. Only two studies, so far, have shown a possible link between IRF-5 and the lipid metabolism. The first one by Montilla *et al.* suggests that IRF-5 may be involved in the lipid metabolism, because *Irf5^-/-^* mice do not properly process myelin-derived lipids in a model for experimental autoimmune encephalomyelitis (EAE) (60). The second study by Orliaguet *et al* demonstrates that IRF-5-deficient macrophages have a highly oxidative nature in the context of a short-term high fat diet. This was caused by transcriptional de-repression of the mitochondrial matrix component *Ghitm*, which is a direct target of IRF-5 (55). GHITM is an inner membrane protein that maintains mitochondrial architecture for efficient oxidative phosphorylation (61) and de-repression of *Ghitm* in *Irf5^-/-^* macrophages results in an increased oxygen consumption rate (55). In our model, IRF-5-deficient p14 CD8 T cell do not display a higher oxygen consumption rate upon *in vitro* stimulation with the gp33 peptide. On the contrary, the cellular respiration was dramatically impaired. Our transcriptomic analysis revealed that several of the downregulated genes in *Irf5^-/-^* CD8 T cells are involved in lipid synthesis (e.g., *Awat2*, *Gpat2*), electron transport chain (e.g., *Coq4*) or in the assembly of the mitochondrial respiratory chain complex I (e.g., *Ndufaf3*, *Ndufb4*). Remarkably, all downregulated genes were directly of indirectly involved in the mitochondrial oxidative phosphorylation process or were components of the mitochondrial envelop. We also found higher *Ghitm* mRNA levels in *Irf5^-/-^*compared with WT p14 CD8 T cells, suggesting that de-repression and rewiring towards oxidative phosphorylation may also occur in CD8 T cells. Nevertheless, *Irf5^-/-^* CD8 T cells were still unable to gain energy from the lipid metabolism. Interestingly, some of the genes that were downregulated in the absence of IRF-5, like *Ndufaf3* and *Coq4*, are associated with mitochondrial disorders in humans (52, 53, 62) and with the Leigh syndrome (58). In our model, the membrane potential of *Irf5^-/-^* p14 CD8 T cells was significantly reduced when compared with WT p14 CD8 T cells at d15 p.i., suggesting that mitochondria are also defective in murine IRF-5 deficient cells. Further investigations are warranted to identify upstream signals and downstream targets of IRF-5 and understand the relationship between IRF-5 and PD-1 and Blimp-1, two molecules involved in metabolic rewiring and mitochondrial restructuring in exhausted cells (9, 10, 11); to note, Blimp-1 is a known target of IRF-5 (63).

IRF-5 deficiency in CD8 T cells was also accompanied by a severe defect in energy production via glycolysis. These results were not entirely unexpected, as Hedl *et al.* reported a role for IRF-5 in enhancing glycolysis in macrophages upon pattern recognition receptor (PRR) stimulation (44). IRF-5 was also shown to regulate glycolysis upon TLR3 stimulation of airway macrophages by directly controlling *Hk2* expression (45). Interestingly, despite displaying a lower ECAR rate upon gp33 stimulation, *Irf5^-/-^*p14 CD8 T cells upregulated several genes involved in the carbohydrate metabolism. This heightened expression of glycolytic genes could represent a compensatory mechanism to cope with defective lipid metabolism or derive from a skewed balance between IRF-4 and IRF-5. Although we did not see any difference in the level of IRF-4 expression (mRNA and protein) at any time point after LCMV Cl13 infection, these two transcription factors are known to compete with each other (41). IRF-4 was also shown to regulate glycolysis in CD8 T cells and to promote expansion and differentiation into effector cells (64); not surprisingly, it also promotes exhaustion (26). IRF-4 is downstream of TCR signaling and links antigen affinity with the cellular metabolism. Indeed, high affinity peptides induce glycolysis and promote differentiation into effector cells via IRF-4, which directly targets *Hk2* (encoding for Hexokinase 2; also an IRF-5 target (45)) and *Glut3* (encoding for a glucose transporter) (64). Thus, it is possible that the higher frequency of KLRG1^+^ IRF-5-deficient p14 cells compared to WT p14 CD8 T cells observed at d8 p.i. derives not only from heightened proliferation, but also from a skewed IRF-4/IRF-5 balance, where IRF-4 is left without a competitor. IRF-4 was also shown to regulate *Hif1a* (64). The Hypoxia Inducible Factor −1ɑ (HIF-1ɑ) itself also promotes glucose metabolism and the expression of glycolytic genes in CD8 T cells (65). Hence, it is possible that HIF-1ɑ and IRF-4 work together to antagonize IRF-5 during clonal expansion promoting glycolysis and limiting the cell division brake imposed by IRF-5. This would be in agreement with a recent study that demonstrates how HIF-1ɑ directly suppresses IRF-5 in monocytes (66). Like IRF-4, hypoxia also promotes functional exhaustion (67). Nevertheless, a skewed IRF-4/RF-5 balance cannot entirely explain the severe defects we observed in IRF-5-deficient CD8 T cells during the chronic phase of infection. Interestingly, the absence of IRF-5 mostly impacted CD8 T cell responses during the chronic phase of infection, when IRF-5 seems to contribute to rewiring the cellular metabolism and helping CD8 T cells cope with the hostile environment created by the chronic inflammation and the continuous antigen stimulation. Even though some glycolytic genes were upregulated in IRF-5-deficient CD8 T cells, these cells were less capable of producing ATP from glycolysis, suggesting that the IRF-5’ role in remodelling the mitochondrial envelope may be crucial for gaining energy from both glycolysis and lipid metabolism and ultimately for cell survival during chronic infections.

IRF-5 deficiency in CD8 T cells lead to premature cell death. In these cells, cell death could have occurred because of exhaustion or the severe bioenergetic defects, or, more importantly, because of lipid peroxidation. Lipid peroxidation is a well-known cause of cell death and typically leads to membrane rupture (68). Moreover, lipid peroxidation can induce apoptosis (69), necroptosis (70), pyroptosis (71), or be the primary driver of ferroptosis (72). Future studies will investigate the cause of death in IRF-5-deficient CD8 T cells and the possible role of IRF-5 in limiting oxidative stress.

In conclusion, we have identified a role for IRF-5 in protecting CD8 T cells from premature functional exhaustion and cell death by regulating the cell cycle and contributing to the metabolic rewiring during chronic LCMV Cl13 infection. Because of the increase in lipid peroxidation in *Irf5^-/-^* CD8 T cells, we believe that IRF-5 could also be involved in protecting CD8 T cells against oxidative stress at chronic stage of LCMV Cl13 infection.

## Materials and Methods

### Mice

C56BL/6-CD45.1 and *Irf5^flox/flox^* mice were purchased from The Jackson Laboratory. *Irf5^-/-^* mice were generated by breeding *Irf5^flox/flox^* mice with mice expressing *Cre*-recombinase under the CMV promoter. Mice with a targeted IRF-5 mutation in T cells were generated by breeding *Irf5*^flox/flox^ mice with mice expressing the *Cre*-recombinase under the Lck promoter (38). LCMV gp33 antigen-specific p14 mice were crossed with *Irf5*^flox/flox^ mice to obtain *Irf5 ^lox/flox^* p14 mice. These mice were crossed with *Irf5^-/-^* mice to obtain IRF-5 deficient LCMV gp33-antigen specific p14 mice (*Irf5^-/-^* p14). All mice were housed and bred at the INRS animal facility under specific pathogen-free conditions and were used at 6-10 weeks of age.

### Ethical statement

*In vivo* experiments were performed under protocols approved by the Comité Institutionnel de Protection des Animaux of the INRS - Centre Armand Frappier (#1910-01, #2003-02). These protocols respect procedure on good animal practice provided by the Canadian Council on Animal Care.

### Viral production and titration

LCMV clone 13 was originally obtained from Dr. Sam Basta (Queens University) and expanded on L-929 cells in EMEM (Wisent) supplemented with 1mM Sodium Pyruvate (Wisent) and 5% FCS (PAA). Cells were infected at a MOI of 0.02 and supernatants collected and cleared of debris 48 h after infection. Viral titers of viral stocks and processed organs were determined on MC57 cells using a standard LCMV focus-forming assay (73).

### Peptide and tetramers

The synthetic peptide gp_33–41_: KAVYNFATC (LCMV-GP, H-2Db) was purchased from New England Peptide (now Biosynth; Gardner, MA, USA). PE-gp_33–41_ tetrameric complexes were synthesized in-house and used to detect LCMV-specific CD8 T cells (74). Briefly, splenocytes were first stained with PE-gp_33–41_ tetramers for 30 minutes at 37°C, followed by direct surface staining with the described antibodies for another 20 minutes on ice (75).

### Cell isolation and adoptive transfer experiments

LCMV- specific CD8 p14 T cells were isolated using negative selection (Miltenyi Biotec) from spleens of *Irf5^flox/flox^* - *CMV-Cre^neg^*-p14 (WT p14) and *Irf5^flox/flox^* – *CMV-Cre^+^-*p14 (*Irf5^-/-^* p14) mice as previously described (76). For adoptive transfer experiments, cells were stained with anti CD3-Alexa fluor 700 (500A2, eBioscience), anti CD8-PE (53-6.7, BD Bioscience), anti CD44-PE-Cy7 (IM7, BD Bioscience), anti CD62L-APC (MEL-14, BD Bioscience), and naïve cells were sorted based on their CD62L^high^ CD44 ^low^ phenotype. 2000 cells were transferred intravenously into congenic CD45.1 mice 1-day prior infection with 2 x10^6^ PFU LCMV clone 13.

For RNA sequencing and digital droplet PCR (ddPCR), adoptively transferred p14 cells was sorted 21 days post infection. For RNA sequencing experiment, 4 samples of 4 individual experiments including 2 males and 2 females per group were used. 8 WT males and 10 KO males were pooled for the 1^st^ experiment, 8 WT males and 17 KO males for the 2^nd^ experiment, 7 WT females and 17 KO females for the 3^rd^ experiment, and 8 WT females and 17 KO females for the 4^th^ experiment.

For ddPCR experiment, 3 samples of 3 experiments using females were used. Each sample is the pool of isolated cells from 10-15 spleens. Cells were stained with anti CD3-Alexa fluor 700 (500A2, eBioscience), anti CD8-PE (53-6.7, BD Bioscience), anti CD45.2-FITC (104, BD Bioscience). Cells were sorted on a BD FACSAria IIu cell sorter from BD Biosciences, 3 laser configurations (Blue 488 nm, Red 633 nm, Violet 405 nm), nozzle size 70-micron, sheath pressure 70 psi. Cell sorting were performed at the Flow cytometry platform of the Institute for Research in Immunology and Cancer’s Genomics (Université de Montreal). CD8^+^ CD45.2^+^ population was collected directly into 500µl Trizol (Sigma) and stored at −80°C until used.

### Flow cytometry

Surface and intracellular staining was performed as previously described (76, 77). The following fluorochrome-conjugated antibodies were used for surface staining anti CD3-Alexa fluor 700 (500A2, eBioscience), anti CD8-PB, aniti CD8-PE, anti CD8-APC (53-6.7, BD Bioscience), anti CD45.2-FITC (104, BD Bioscience), anti KLRG1-APC (2F1, eBioscience), anti CD127-PE (SB/199, BD Bioscience), anti LAG-3-PE-Cy7 (C9B7W, eBioscience), anti PD-1-biotin (29F.1A12, Biolegend), anti TIM-3-BV605 (RMT3-23, Biolegend), anti CD44-PE-Cy7 (IM7, BD Biosciences), anti CX3CR1-PerCP-Cy5.5 (SA011F11, Biolegend), anti CD101-Alexa flour 700 (Moushi101, eBioscience), and streptavidin-V500 (BD Bioscience). Cells were fixed using 2% paraformaldehyde and permeabilized using 0.1% saponin solution. Expression of Tibet, EOMOES, and IRF-4 was measured by staining cells with anti-Tbet-PerCP-Cy5.5 (4B10, eBioscience), anti-EOMES-PE (Dan11mag, eBioscience), anti-IRF4-PECy-7 (3E4, eBioscience). For cell cycle analysis, cells were stained using anti Ki67-PE-Cy7 (SolA15, eBioscience), washed with PBS, and labelled with DAPI (10 µg/ml final concentration) at room temperature for 10 minutes. To measure IRF-5 expression, permeabilized cells were stained with anti IRF5-PE (W16007B, Biolegend) and analyzed by flow cytometry or then stained with DAPI and acquired by ImageStreamX as previously described (38).

For cytokine quantification, splenocytes were restimulated for 5 hours at 37°C with 1 μg/ml of MHC class I-restricted LCMV peptide gp_33-41_ in the presence of 2 ng/ml recombinant human IL-2 (Peprotech), 10% fetal bovine serum (premium FBS, Wisent Bioproduct), and 1 µg/ml Brefeldin A (BD Bioscience). Following *ex vivo* restimulation, cells were stained with stained, fixed, and permeabilized as described above. The following antibodies were used for intracellular staining: anti IFNγ-APC (XMG1.2, BD Biosciences), anti TNF-PE-Cy7 (MP6-XT22, BD Biosciences), and anti IL2-PE (JES6-5H4, eBioscience). Samples were acquired on a BD LSRFortessa II cell analyzer (Becton Dickinson). Data was analyzed with Flowjo v10.8.1.

### Bulk RNA sequencing

Total RNA was isolated using RNeasy mini kit (Qiagen) according to the manufacter’s instructions. RNA was quantified using Qubit (Thermo Scientific) and quality was assessed with the 2100 Bioanalyzer (Agilent Technologies). Transcriptome libraries were generated using the KAPA RNA HyperPrep (Roche) using a poly-A selection (Thermo Scientific). **S**equencing was performed on the Illumina NextSeq500, obtaining around 20M reads per sample. Bulk RNA sequencing was performed at the Genomics Platform of the Institute for Research in Immunology and Cancer (Université de Montreal).

### RNA seq analysis

Adapter sequences and low-quality bases in the resulting FASTQ files were trimmed from raw sequences using Trimmomatic version 0.35 (78) and genome alignments were conducted using STAR version 2.7.1a (79). The sequences were aligned to the mouse genome version GRCm38, with gene annotations from Gencode version M25 based on Ensembl 100. As part of quality control, the sequences were aligned to several different genomes to verify that there was no sample contamination. Expression levels were obtained as gene readcounts from STAR as well as estimated using RSEM (80) in order to obtain gene and transcript level expression in reads per transcripts per million (TPM) for these stranded RNA libraries.

DESeq2 version 1.30.1 (81) was then used to normalize gene readcounts and compute differential expression between the various experimental conditions. Significant gene sets were chosen according to a false discovery rate (FDR) cut-off of 5% (padj < 0.05).

Computing both a hierarchical sample clustering based on Pearson correlation of normalized log readcounts and a principal component analysis shows that samples behave as expected, with no obvious outlier, as well as good consistency among replicates.

In addition to the various pairwise comparisons performed, a multivariate model was also built in DESeq2, and a likelihood ration test (LRT) used to isolate the treatment effect from contributions of the other factors in play, namely sex and time. RNAseq analysis was performed at the Genomics Platform of the Institute for Research in Immunology and Cancer (Université de Montreal).

### Gene ontology analysis

Up-regulated and down-regulated genes were determined with the cut-off SCORE of ≥ |0.7| in the Rank gene list (table 1). These genes were compared using Gene Ontology analysis; biological processes associated with a p < 0.05 were considered as significantly enriched. Genes were compared with gene sets of different biological functions. A cut-off of fold change ≥ |2| was used to determine differentially regulated genes. Heatmaps were generated based on the readcount normalization using Usegalaxy.org.

**Table 1:**
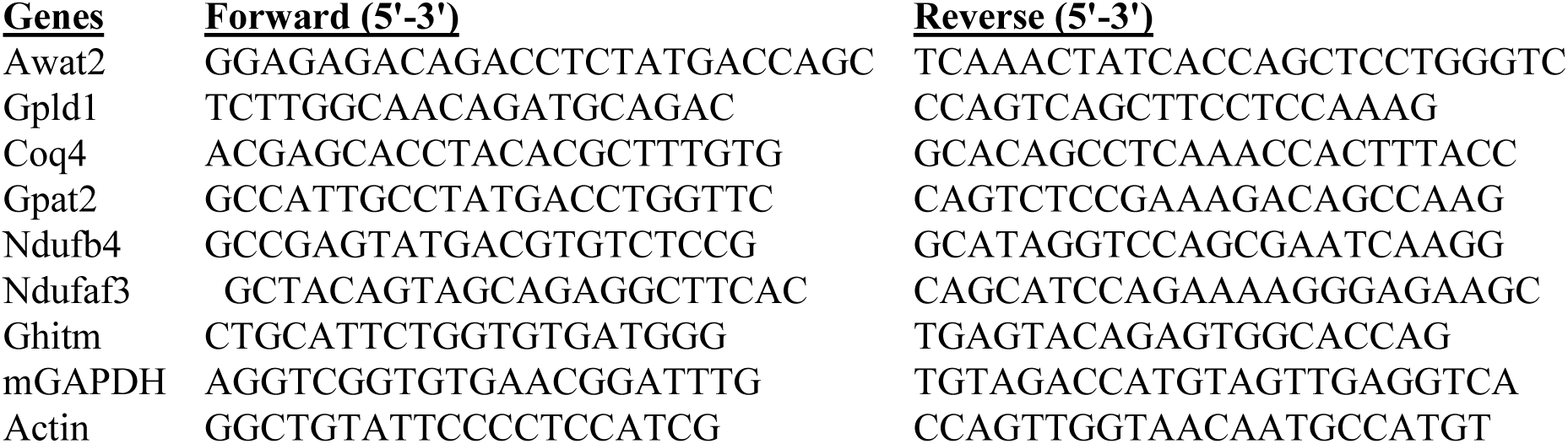
Primers used to determine relative gene expression by ddPCR.

### Metabolic assays

CD8 T cells from WT p14 or *Irf5^-/-^* p14 mice were isolated as described above and cultured with or without 1µg/ml gp33 peptide in complete RPMI medium supplemented with 2ng/ml recombinant human IL-2 (Peprotech). Cells were then collected at 6 or 24 hours after incubation and washed with PBS. For mito stress and real-time ATP production experiments, 4 replicates per condition were used and 2 x 10^5^ cells were plated for each replicate. Cells were resuspended in Agilent RPMI media containing 10 mM glucose, 2 mM glutamine, and 1 mM sodium pyruvate and incubated at 37° C for 1 h without CO_2_. Mitochondria respiration, oxygen consumption rates (OCR), and extracellular acidification rates (ECAR) were measured using the XF-96 Extracellular Flux Analyzer (Agilent) at baseline and in response to 1.5 µM Oligomycin, 2 µM FCCP, and 0.5 µM rotenone/antimycin A, according to manufacturer’s protocols (Mito Stress Test Kit, Agilent). For real-time ATP production, oxygen consumption rates (OCR) and extracellular acidification rates (ECAR) were measured at baseline and in response to 1.5 µM Oligomycin and 0.5 µM rotenone/antimycin A following manufacturer’s protocols. The results were analyzed using Wave software (Agilent).

To measure glucose and lipid uptake capacity, cells were labelled with 150 μM of the fluorescently labeled glucose analog 2-NBDG (Thermo Fisher Scientific, N13195) or with 1µM bodipy (Thermo Fisher Scientific, D3821) in serum-free media for 30 minutes at 37°C. Cells were washed with PBS prior to cell surface staining, as described above, and analyzed by flow cytometry.

### MitoSox and tetramethylrhodamine methyl ester (TMRM)

To measure mitochondrial superoxide and the mitochondrial membrane potential state, cells were labelled with 5 μM MitoSOX Red mitochondrial superoxide indicator reagent (ThermoFisher) or with 100 nM TMRM (ThermoFisher) and were incubated for 30 min at 37 °C. Cells then were washed with PBS prior to cell surface staining as described above and analyzed by for flow cytometry.

### Lipid peroxidation

To evaluate lipid peroxidation, cells were labelled with Image-iT® Lipid Peroxidation Sensor (ThermoFisher) at a final concentration of 5 µM, then were incubated for another 30 minutes at 37°C. Cells were then washed three times with PBS and stained with other cell surface antibodies. The fluorescence signals were read at separate wavelengths by flow cytometry; one at 581/591 nm for the live dye, and the other at 488/510 nm for the oxidized dye. The ratio of the emission fluorescence intensities at 590 nm to 510 nm gives the read-out for lipid peroxidation in cells. Lower ratio corresponds to higher level of lipid peroxidation.

### Digital droplet PCR (ddPCR)

Total RNA was isolated using RNeasy mini kit (Qiagen) according to the manufacturer’s instructions. Reverse transcription was performed using the iScript cDNA synthesis kit (Bio-Rad) following the manufacturer’s protocol. 25 µl reaction mix for ddPCR was made by 12.5 µl EvaGreen supermix, 250 nM forward and reverse primers, 2 µl cDNA and water. Sequences of primers are in Table 1. ddPCR was performed using a QX200 Droplet Digital PCR system (Bio-Rad). Specifically, 20 µl of each reaction mix and 70 µl of oil were converted to droplets with the QX200 droplet generator (Bio-Rad). Droplet-partitioned samples were then transferred to a 96-well plate, sealed, and cycled in a C1000 Touch Thermocycler (Bio-Rad) under the cycling protocol that was optimized for each primer. The cycled plate was then transferred and read in the QX200 reader (Bio-Rad) and analyzed by QX manager software 2.0 (Bio-Rad). Fold change expression relative to naïve was calculated by normalized with *Gapdh* and *Actb*.

### Statistical analysis

GraphPad Prism 8.0 was used for statistical analysis. Each value represents at least three independent experiments. Two-tailed Student t test or ordinary one-way ANOVA or two-way ANOVA were used to determine statistical significance. * denotes p<0.05, ** denotes p<0.01 *** denotes p< 0.001 **** denotes p< 0.0001. Error bars denote ± SEM.

## Supplementary materials

Supplementary Figure 1-7

Fig. S1: Adoptive transfer experiment design and mouse survival rate

Fig. S2: CD8 T cell percentages and numbers in LCMV Cl13-infected *Irf5^f/f^x Lck-Cre^+^*

Fig. S3: Cytokine production by CD8 T cells adoptively transferred in female mice

Fig. S4: Expression of IRF-4, Eomes, and T-bet in adoptively transferred CD8 T cells

Fig. S5: Cytokine production by CD8 T cells and viremia in *Irf5^f/f^x Lck-Cre^+^*mice

Fig. S6: Heatmap of the first 45 differentially expressed genes

Fig. S7: Cellular metabolism analysis of WT and *Irf5^-/-^* p14 CD8 T cells from male mice

Supplementary Table 1: List of the first 45 differentially expressed genes

## Acknowledgements

We thank the Canadian Institutes of Health Research (PJT-190001 to SS.; PJT-148614 to KMH; PJT-175173 to JHF) for financial support. We thank our animal care technician Annie Salesse, who cared for the mice used in this study. LTM was supported by scholarships from the Fondation Armand-Frappier and the Fonds de Recherche du Québec – Santé (FRQS).

## Author contributions

LTM: performed and analyzed experimental data. SSw, THN, TC, LCP: performed experiments; HL provided expertise; AL provided essential materials; KHM and JHF: provided conceptual input. S.S. analyzed experiments and supervised the project. LTM and SS wrote the manuscript with input from all coauthors.

## Competing interests

Authors declare that they have no competing interests.

---

**Supplemental Figure 1.**
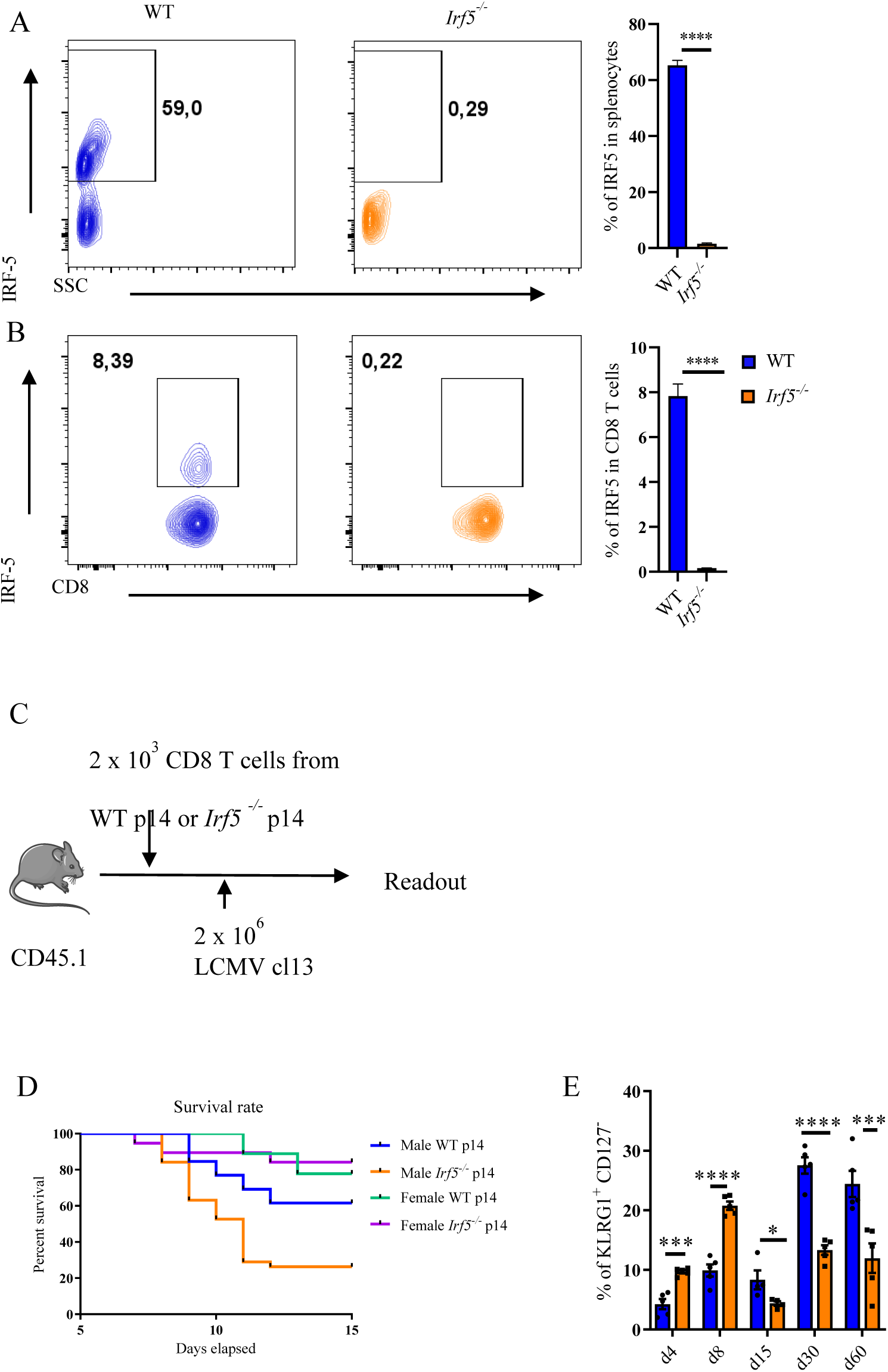
(A-B) Representative FACS plots and graphs representing IRF-5 expression by total splenocyte (A) and CD8 T cells (B) from WT (*Irf5^f/f^ x CMV-cre^-^*) and *Irf5^-/-^* ( *Irf5^f/f^ x CMV-cre^+^*) p14 transgenic mice. (C) Experimental set-up for adoptive transfer experiments described throughout the manuscript. (D) Graph illustrates the survival rate of male or female CD45.1 recipient mice adoptively transferred or not with WT and *Irf5^-/-^* p14 CD8 T cells over the course of LCMV Cl13 infection. (E) Graph shows the percentage of KLRG1^+^ CD127^-^ WT and *Irf5^-/-^* p14 CD8 T cells found in the spleen of recipient female mice over the course of infection

**Supplemental Figure 2.**
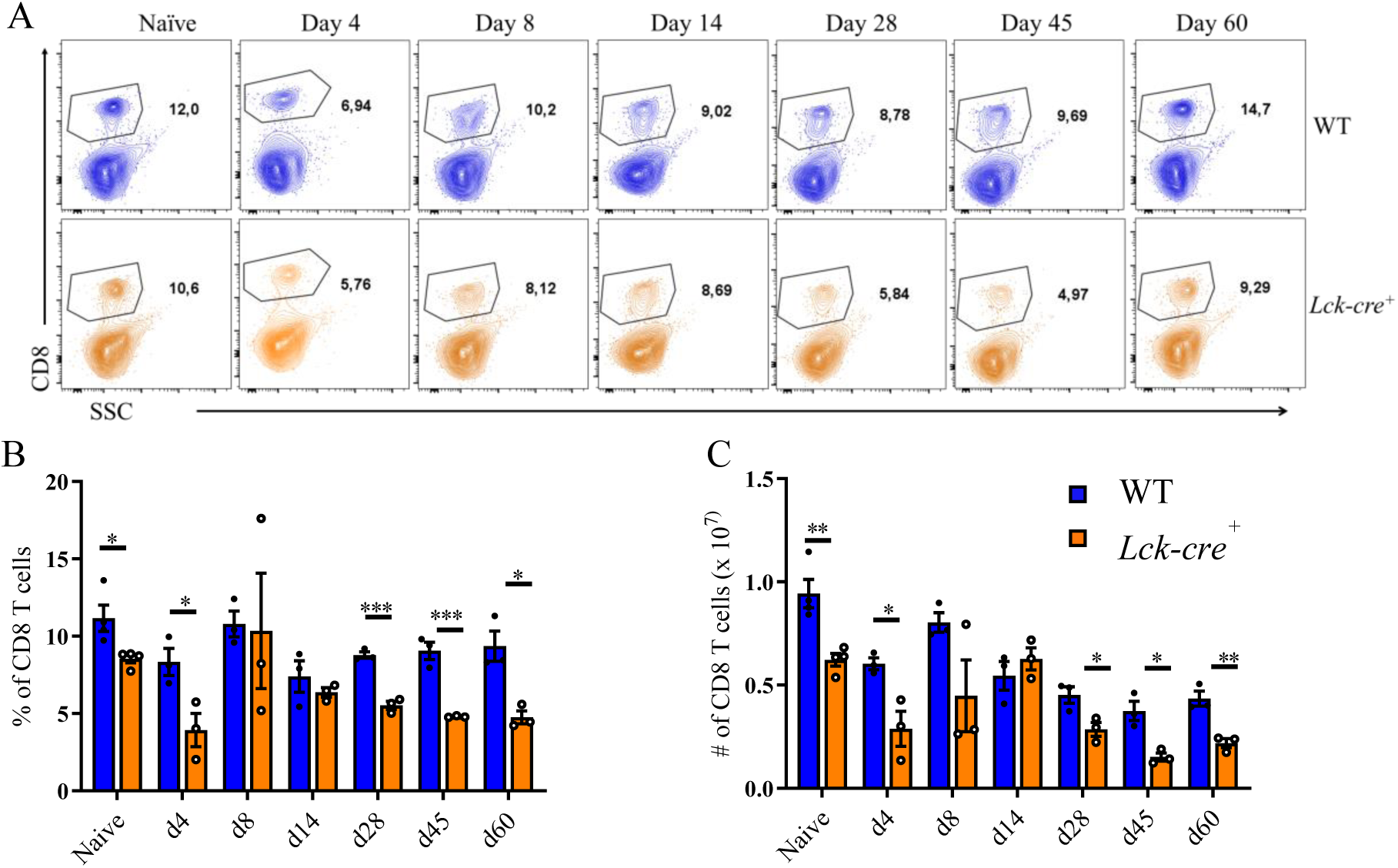
I*r*f5f*^/f^x Lck-Cre^-^* (WT) and *Irf5^f/f^x Lck-Cre^+^* (Lck-cre^+^) mice were infected intravenously with 2 × 10^6^ PFU LCMV Cl13 and euthanized at various time points p.i.. Graphs show (A) representative FACS plots for CD8 T cell staining, and (B) the percentages and (C) absolute numbers of CD8 T cells found in the spleen over the course of infection.

**Supplemental Figure 3.**
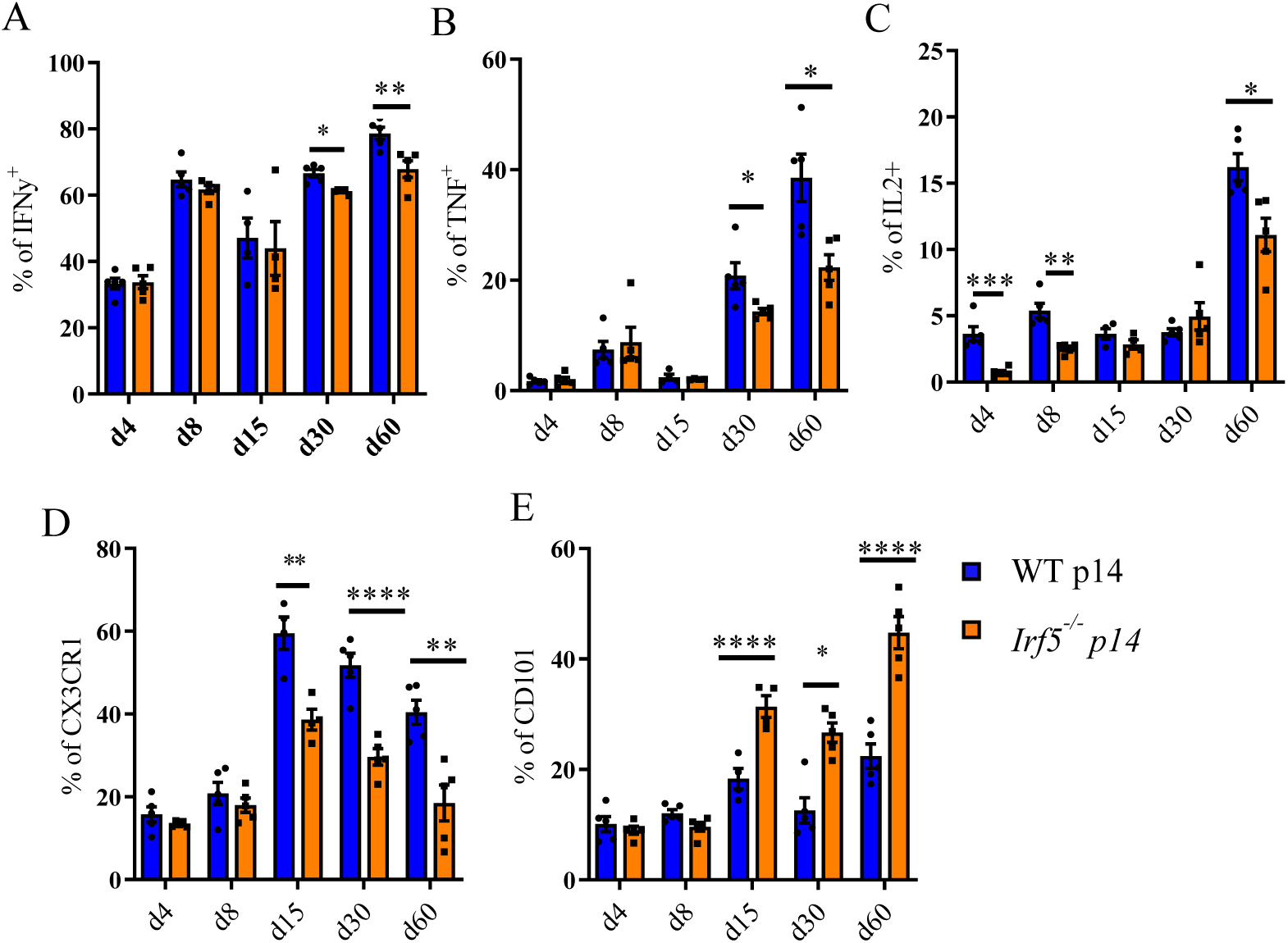
CD8 T cells from female WT or *Irf5^-/-^* p14 mice were adoptively transferred into CD45.1 recipient mice one day prior to intravenous infection with 2 × 10^6^ PFU LCMV Cl13. Mice were euthanized at various time points after infection. Graphs show the frequency of (A) IFNγ^+^, (B) TNF^+^, and (F) IL-2^+^ CD8 T cells upon *in vitro* restimulation with the gp33 peptide; and the percentage of (D) CX3CR1^+^ and (E) CD101^+^ splenic CD8 T cells.

**Supplemental Figure 4.**
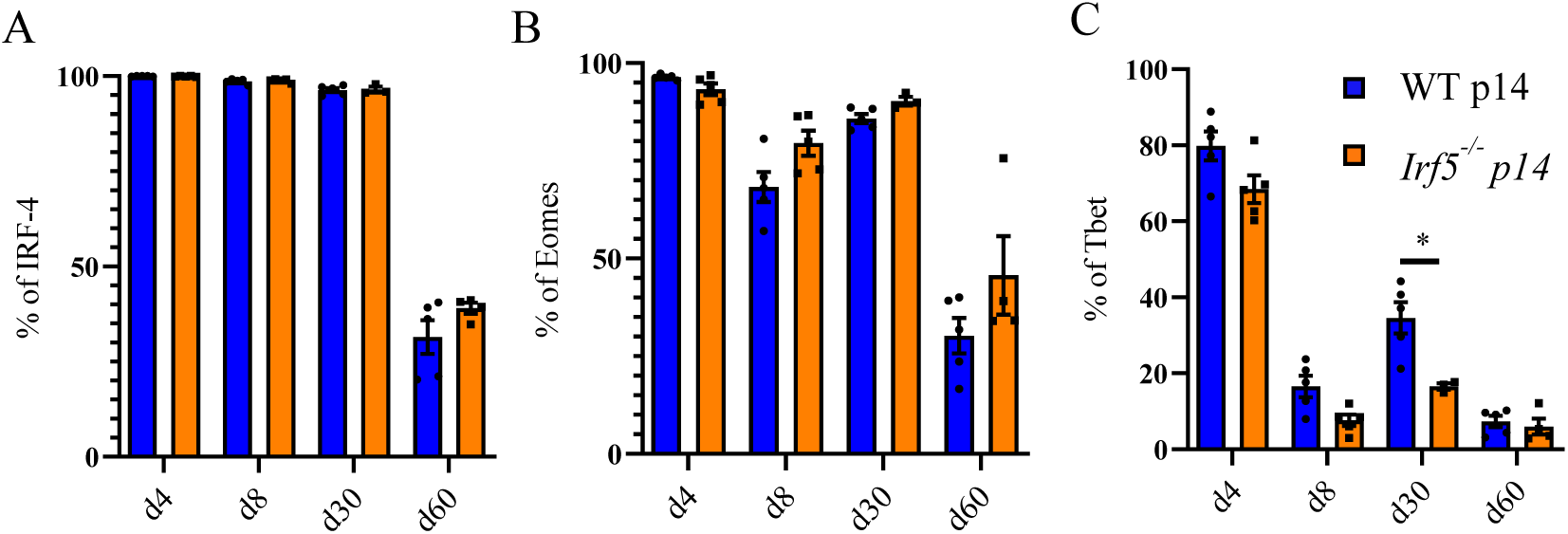
Graphs illustrate the frequency of (A) IRF-4^+^, (B) Eomes^+^, and (C) T-bet^+^ WT and *Irf5^-/-^* p14 CD8 T cells found in the spleen of recipient mice at various time points after LCMV Cl 13 infection.

**Supplemental Figure 5.**
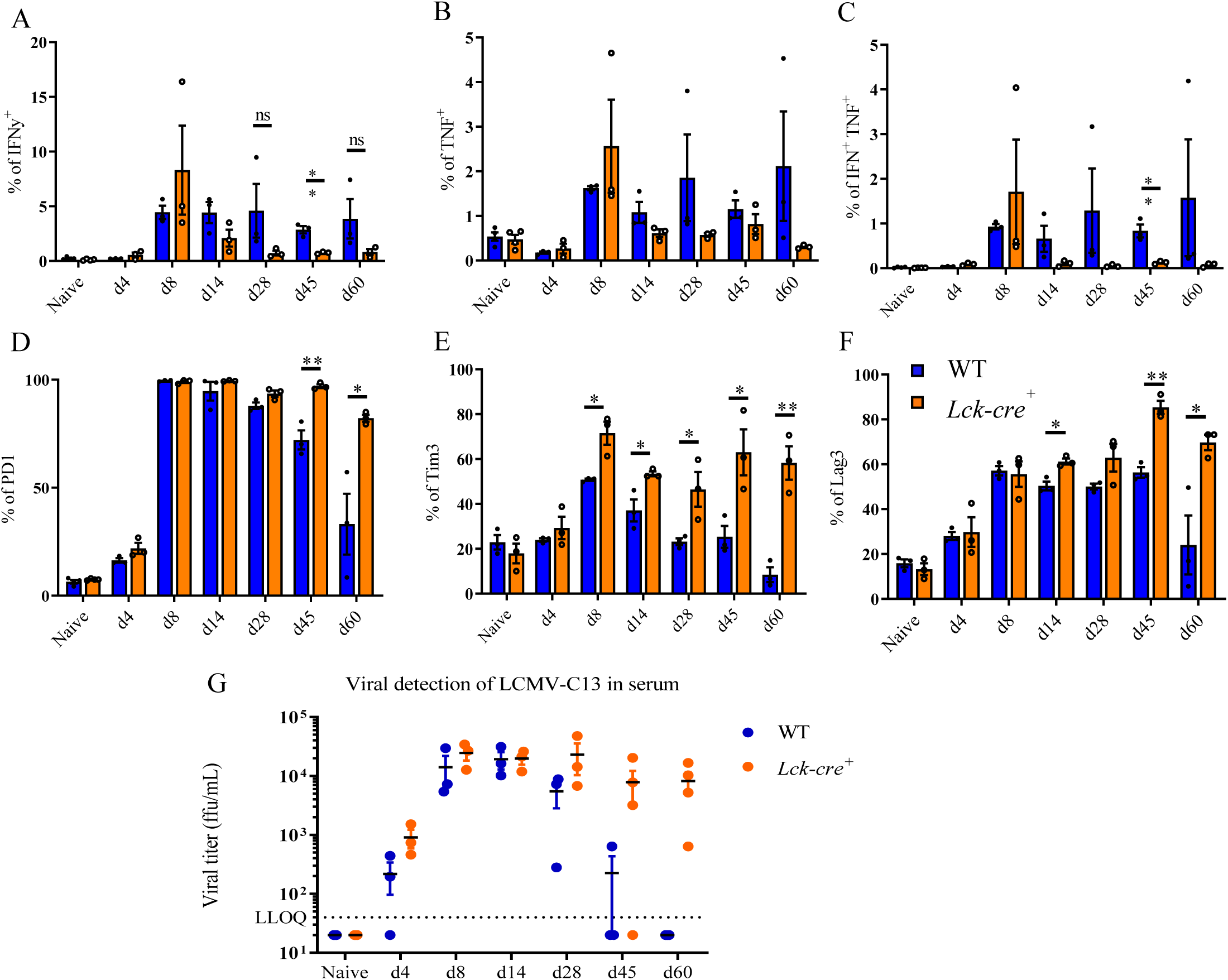
I*r*f5f*^/f^x Lck-Cre^-^* (WT) and *Irf5^f/f^x Lck-Cre^+^* (Lck-cre^+^) mice were infected intravenously with 2 × 10^6^ PFU LCMV Cl13 and euthanized at various time points p.i.. Graphs show the frequency of (A) IFNγ^+^, (B) TNF^+^, and (F) IFNγ^+^ TNF^+^ CD8 T cells upon *ex vivo* restimulation with the gp33 peptide; the percentage of gp33-tetramer^+^ splenic CD8 T cells expressing (D) PD-1, (E) TIM-3, and (F) LAG-3 over the course of infection. (G) Viral titers were assessed in the serum of infected mice until d60 p.i.

**Supplemental Figure 6.**
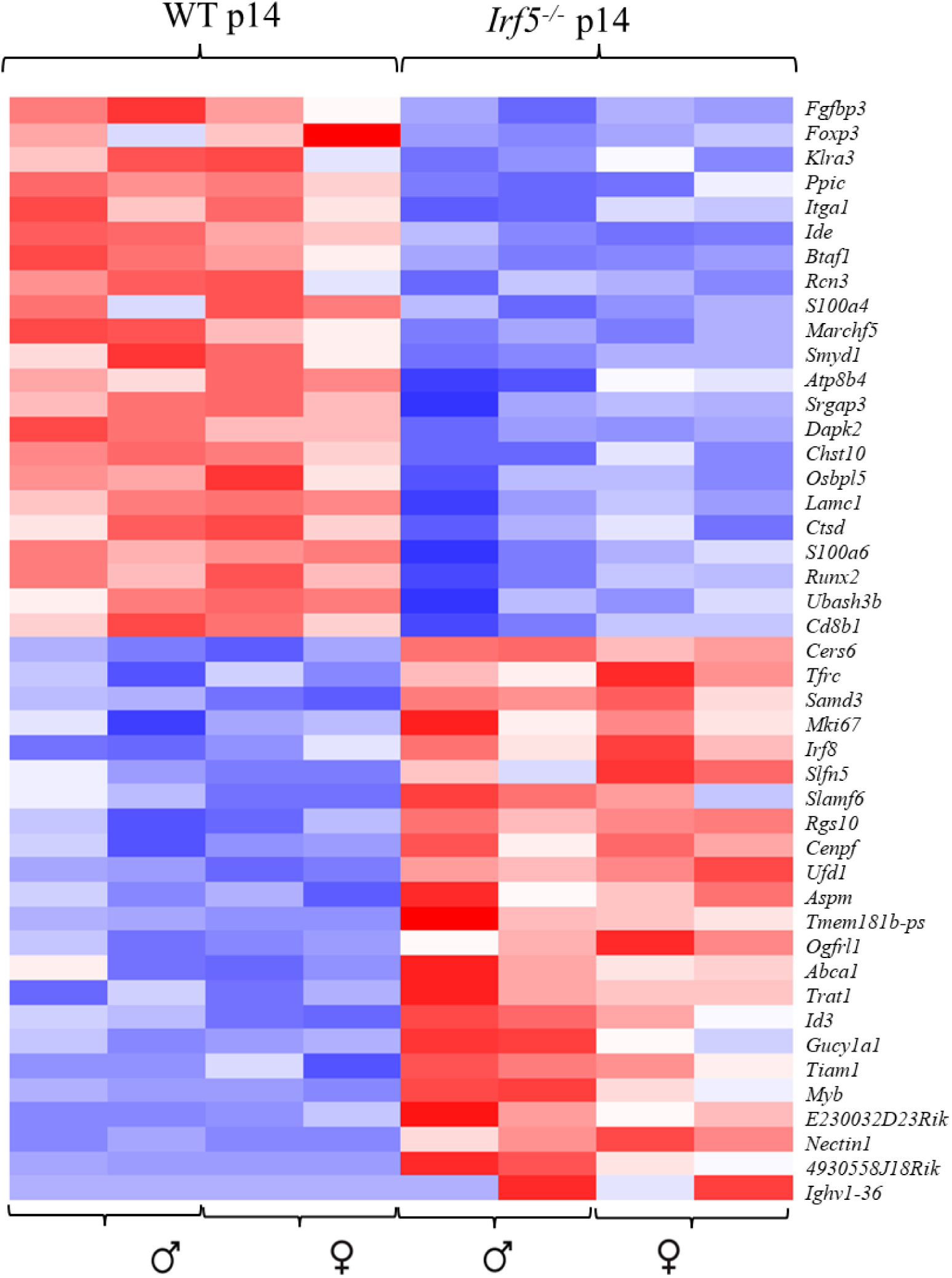
The heatmap shows the top 45 differentially expressed genes with padj values smaller than 0.05) in WT and *Irf5^-/-^* p14 CD8 T cells isolated from female and male recipient mice at d21 p.i..

**Supplemental Figure 7.**
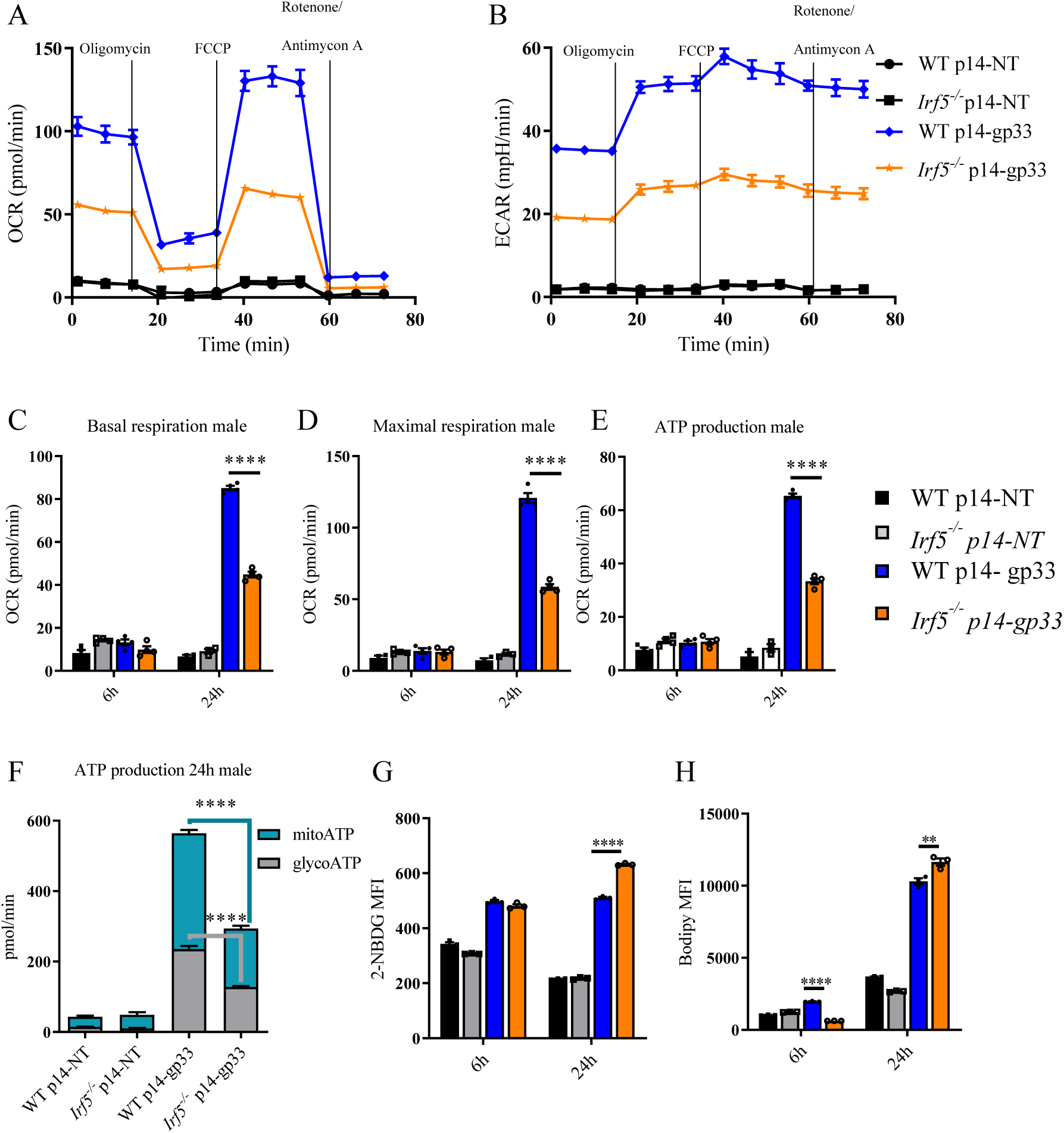
WT and *Irf5^-/-^* p14 CD8 T cells purified from male mice were cultured *in vitro* in the presence or absence of the gp33 peptide, and the mitochondria respiratory capacity was measured using the Seahorse XFe-96 analyzer. Graphs illustrate (A) the representative oxygen consumption rates (OCR) and (B) the extracellular acidification rates (ECAR) over time; (C) the basal and (D) maximal respiration, and (E) the ATP production at 6h and 24h after stimulation with the gp33 peptide; (F) the ATP production rate; and (G) the glucose (measured using the fluorescent glucose analog 2-NBDG) and (H) the fatty acid uptake capacity (quantified using bodipy FLC_16_ fluorescence intensity).

## Notes

### Competing Interest Statement

The authors have declared no competing interest.

